# Hindmarsh–Rose neuronal network with spike-timing-dependent plasticity demonstrates coordinated reset neuromodulation for Parkinson’s disease

**DOI:** 10.64898/2026.05.27.728228

**Authors:** Shahin Sharafi, Jesse I. Gilmer, Mazen Al Borno, Thomas K. Uchida

**Author notes:** Contributing authors.

## Abstract

Computational models of brain structures impacted by Parkinson’s disease are useful for studying pathological synchronization and exploring potential therapies. We use the Hindmarsh–Rose neuronal model to simulate synchronized activity in the subthalamic nucleus, capturing key features of the pathological rhythms associated with parkinsonian tremor. Our model incorporates chemical synapses whose strengths evolve according to a spike-timing-dependent plasticity (STDP) rule. We apply coordinated reset stimulation with rapidly varying sequences (RVS CR) and examine its ability to weaken synaptic weights, thereby reducing neuronal synchrony. This stimulation technique delivers phase-shifted stimuli to distinct sites in a random sequence. We explore how stimulation frequency and the number of stimulation sites affect the efficacy of RVS CR at desynchronizing the network and observe good agreement with previous studies. We demonstrate that RVS CR efficacy is sensitive to the depression-to-potentiation ratio in the STDP rule, which may be an important parameter to tune when reconciling simulations with experimental data. Numerical simulation of neuronal networks is constrained by computational resources when models demand large networks. This work proposes a model that demonstrates similar utility with a relatively small network, enabling researchers to study pathological neuronal activity and treatments more efficiently.

## 1 Introduction

Parkinson’s disease (PD) is a common neurodegenerative disease, the global prevalence of which is second only to Alzheimer’s disease [1]. The number of individuals living with PD is expected to double by 2050 [2]. PD is associated with dopamine depletion in the substantia nigra pars compacta [3, 4], resulting in a significant increase in neuronal synchronization in the subthalamic nucleus (STN) and internal globus pallidus (GPi) [5–8]. Levodopa and other pharmacological treatments can be highly effective at alleviating motor symptoms during the first 5–10 years of PD therapy [9, 10]; however, their effectiveness typically diminishes over time. Additionally, prolonged levodopa treatment can cause side effects such as increasingly frequent and severe involuntary movements (dyskinesias) [11]. Surgery may be considered as an alternative treatment for individuals with PD when medication is no longer effective at managing symptoms. Lesion surgery involves selectively destroying brain tissue to interrupt the connections between the globus pallidus and the thalamus [12]; however, this treatment has been largely replaced by deep brain stimulation, which is reversible and mimics the effect of tissue lesioning but with fewer side effects [13, 14].

Deep brain stimulation (DBS) has proven to be an effective therapy for individuals with PD who have drug-resistant motor symptoms [15]. In high-frequency DBS therapy, periodic high-frequency electrical pulses (over 100 Hz) are delivered to the brain tissue via surgically implanted electrodes. Over the past decade, the STN (a small subcortical nucleus comprising approximately 550,000 neurons [16]) has been recognized as an effective target for the placement of DBS electrodes to suppress excessive neuronal synchrony [17–19]. High-frequency DBS can significantly reduce the need for medication in treatment of drug-resistant tremor and dyskinesias; however, axial symptoms such as dysarthria, gait disorders, and postural instability are less likely to improve with DBS [20–23]. Moreover, symptoms often reappear soon after the cessation of stimulation [24]; thus, DBS is typically delivered continuously [25]. Side effects of DBS include dysarthria (impaired speech and reduced verbal fluency), dysesthesia (unusual touch-based sensation), and ataxia (poor balance and unsteady gait) [26, 27]. Additionally, patients must be in good overall health to be candidates for the invasive and lengthy surgery to implant the electrodes [23], and periodic surgeries are required to replace the battery in the internal pulse generator [28]. Unlike periodic DBS techniques, the open-loop coordinated reset (CR) technique is applied by dividing the neuronal population into subpopulations and resetting the phases of the subpopulations so that their spiking activity is evenly spaced over time [27]. The open-loop CR technique is robust to variations in system parameters [27, 29] and has been shown, both theoretically and clinically, to reduce pathological neuronal synchronization in the STN [30] with long-lasting effects [25]. The stimulation used in the open-loop CR technique typically comprises a high-amplitude pulse followed by a low-amplitude pulse: the former aims to “reset” a cluster of neurons, shifting it from a stable synchronized state to a desynchronized state in which the phases of its neurons are uniformly distributed over each period; the latter is applied following a delay and aims to reinforce full desynchronization of the system [27, 29, 31]. Effective cluster resetting requires precise calibration of stimulation parameters (e.g., the delay between the high- and low-amplitude pulses) and even slight variations in these critical parameters can reduce effectiveness [32–34]. Other stimulation techniques have also been investigated in recent studies [35, 36].

Experimental studies suggest that the STN exhibits spike-timing-dependent plasticity (STDP) [37], meaning that the strength of the synapses evolves based on the spike timing of the presynaptic (upstream or proximal) and postsynaptic (downstream or distal) neurons [38]. Pathological neuronal activity associated with PD is characterized by abnormal synchronization and an oscillatory firing pattern, where synaptic weights increase due to the STDP rule [39, 40]. In some model-based studies, incorporating the STDP rule into the system equations has resulted in networks with two possible stable states: one stable synchronized state and one stable desynchronized state [40, 41]. In general, the objective of CR stimulation techniques is to weaken the synaptic weights, thereby reducing neuronal synchrony [39]. Previous modelling studies have shown that the CR technique can move a stable synchronized network (with a high average synaptic weight) to a stable desynchronized state (with a lower average synaptic weight) during application of the CR stimulation and achieve long-lasting neuronal desynchronization [41].

Computational models have been used to explore the effect of CR stimulation parameters on neuronal desynchronization in the STN. Ebert et al. [39] constructed a large-scale computational model of the STN and the external globus pallidus (GPe; 10,000 neurons for each structure) with neuronal connections governed by the STDP rule. They used a single-compartment, conductance-based model to represent the activity of the neurons in the GPe and STN, and demonstrated the efficacy of the CR technique at desynchronizing strongly synchronized neuronal activity. The sensitivity of efficacy to electrode positioning was also demonstrated. Fan and Wang [42] used the Hodgkin–Huxley neuronal model to demonstrate the effectiveness of the triple-structure CR stimulation strategy, where the GPe, GPi, and STN are stimulated simultaneously via three micro-electrodes. Manos et al. [43] investigated the long-lasting effects of CR stimulation in networks modelled using the Hodgkin– Huxley neuronal model with the STDP rule, concluding that acute desynchronization during application of CR stimulation does not necessarily result in long-lasting desynchronization after stimulation ceases for certain stimulation parameters.

In simulation-based studies of neuronal dynamics, computational cost is an important consideration. The Hodgkin–Huxley neuronal model and its descendants comprise equations with dozens of parameters and have high computational expense [44, 45].

Existing studies on how CR stimulation affects neuronal activity have primarily used the leaky integrate-and-fire (LIF) model with the STDP rule. The LIF model is a simplified threshold-based neuronal model with relatively low computational expense; however, it cannot reproduce realistic spiking waveforms or complex spiking patterns [44]. Kromer et al. [25] used the LIF model with STDP to examine how the number of stimulation sites and the stimulation frequency affect long-lasting desynchronization induced by CR stimulation, revealing a nonlinear relationship between these stimulation parameters and efficacy. Khaledi-Nasab et al. [40] studied the effect of CR stimulation parameters on an LIF network of 1,000 neurons with STDP. They observed that randomizing the CR stimulation amplitude, timing, or both enhances the robustness of the long-lasting desynchronization effect. Kromer and Tass [46] performed numerical simulations to investigate the effects of the synaptic connection length scale on the ability of various CR stimulation strategies to induce long-lasting desynchronization. The synaptic connection length scale is a parameter that controls the network architecture—specifically, the degree of local and global connectivity in the network. Kromer and Tass found that CR stimulation parameters can be adapted to the network architecture to enhance the long-lasting desynchronization effect. Kubota and Rubin [47] constructed an LIF network of 100 neurons and used optimization to identify the energetically optimal electric-current waveform for the CR stimulation signal. Although the model of Kubota and Rubin ignored synaptic plasticity, they demonstrated that the effectiveness of CR stimulation can depend on several model and stimulation parameters. Other studies that employed models other than the LIF model did not incorporate the STDP rule in their system equations. Liu et al. [48] used the Izhikevich model to construct a neuronal network of the basal ganglia and thalamus. They applied intermittent noisy stimulation (referred to as coordinated reset noisy stimulation, CRNS) to the GPe, GPi, and STN, and found that CRNS is more effective at suppressing pathological synchronization while consuming less energy; however, synaptic plasticity was ignored for simplicity.

The Hindmarsh–Rose (HR) neuronal model comprises three first-order differential equations and can mimic diverse neuronal behaviours, including chaotic dynamics, while remaining computationally efficient [44, 49, 50]. Jalili [51] used the HR model with constant synaptic weights to study the effects of time delays in the chemical neuronal coupling dynamics, concluding that time delays enhanced phase synchronization in the network. In a subsequent study, Jalili investigated the influence of STDP on a network of 5 HR neurons and concluded that STDP can increase the level of synchrony [52]. Chakravartula et al. [53] constructed a network of 100 HR neurons and introduced a new type of adaptive coupling where the connection strength between neurons depends on their membrane potentials. They demonstrated that their model can exhibit both transient and permanent synchronization within local clusters of neurons. Baran et al. [54] studied the similarity between the STDP rule and the dynamics of a memristor using two coupled HR neurons with both chemical coupling and memristor-based unidirectional synapses. Lysyansky et al. [55] used the HR model to study a large, initially silent neuronal network (i.e., a network in which the synapses do not trigger action potential firing in the postsynaptic neurons [56]) and found that CR stimulation can be effectively applied in such a network. Again, synaptic plasticity was ignored for simplicity.

In this study, we model a network of 100 identical HR neurons with synaptic weights governed by the STDP rule. We compare the effect of two commonly used types of noise on network dynamics and investigate the efficacy of CR stimulation on reducing synchronized neuronal activity associated with parkinsonian tremor. To the best of our knowledge, this is the first study to use an HR model with STDP for examining the effects of noise and the efficacy of CR stimulation. We consider unidirectional excitatory chemical coupling between neurons and tune the HR model and chemical coupling parameters to replicate the behaviour of individual STN neurons as well as population-level dynamics of the STN reported in previous studies [25, 57– 59]. We demonstrate that implementing white noise in the HR model yields a single stable network state of synchronized activity whereas Poisson noise can produce two stable states, one synchronized and one desynchronized. We demonstrate the utility of the proposed model by investigating the effect of the number of stimulation sites and the stimulation frequency on CR stimulation efficacy. The proposed HR-with-STDP model predicts network behaviour that compares favourably with the results from larger models while retaining nonlinear neuronal dynamics, enabling researchers to efficiently explore a wide range of stimulation parameters with a reduced network size.

## 2 Methods

In this section, we describe the HR model of neuronal dynamics, the STDP rule, two types of noise, and the coordinated reset stimulation with rapidly varying sequences (RVS CR) strategy we employ. We also describe the network topology we adopt as well as the concept of the order parameter, which we use to quantify network synchrony.

### 2.1 Mathematical model

The spiking dynamics of neurons in the STN are modelled using the HR neuronal model, with the STDP rule governing the evolution of the synaptic weights. The HR model comprises three first-order differential equations per neuron [49]:

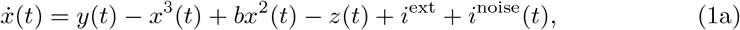

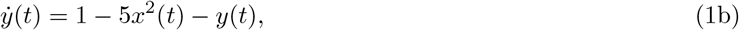

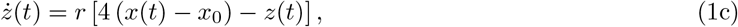

where *x*(*t*) is a dimensionless form of the neuronal membrane potential, *y*(*t*) is a state representing the aggregate effects of the fast current dynamics, and *z*(*t*) is a state representing the aggregate effects of the slow current dynamics. The model includes four constant parameters: *b* regulates the neuron’s dynamics (spiking or bursting), *i*^ext^ is the external input current injected into the neuron, *r* regulates the spiking frequency, and *x*_0_ represents the resting potential. We set parameters *b* = 3.0 and *x*_0_ = *−*1.6, adopting typical values used in the literature [53]; to replicate the neuronal spiking behaviour observed in the STN, we set *i*^ext^ = 1.45 and *r* = 0.01 [57, 59]. In this work, we explore two models for the time-varying noise *i*^noise^(*t*) to compare their effects on the network’s stability characteristics. The first noise model is simple white noise, where *i*^noise^(*t*) is assigned a random value at each time step that is selected from a uniform distribution over the range [−0.5, 0.5] (Supp. Fig. 1). This model assumes that the input received from all unmodelled STN neurons, once these inputs have been scaled by their respective synaptic weights and summed, resembles white noise. The second noise model we explore assumes the unmodelled STN neurons provide an input *i*^noise^(*t*) that resembles an independent Poisson process, following the conductance noise model presented by Kromer et al. [25], with firing rate 50 Hz, noise intensity 3.0, and synaptic timescale 2 ms (Supp. Fig. 2). We generate Poisson spike trains following the method described by Heeger [60]. In all simulations, the initial conditions for the state variables are selected randomly from uniform distributions with the following ranges: *x*(0) [1.5, 1.0], *y*(0) [2.0, 2.0], and *z*(0) ∈ [0, 2.6].

Several studies have assumed that the neurons in the STN are locally connected through unidirectional excitatory chemical synapses, an assumption that is supported by considerable experimental and theoretical evidence [25, 61–63]. Therefore, to model the neuronal dynamics in the STN, we consider a network of *N* = 100 identical HR neurons coupled through unidirectional excitatory chemical synapses:

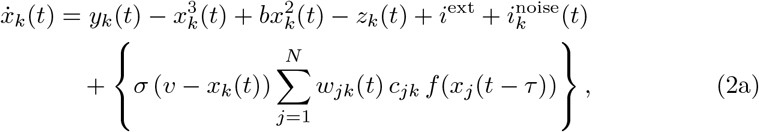

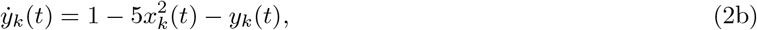

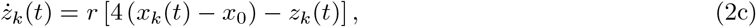

where the braced expression in Eq. (2a) is the fast-threshold-modulation model, which models the effect of the excitatory chemical synapses [64]. The scaling factor *σ* prevents excessive inputs from being injected through the synapses; selection of the value for this parameter is discussed in detail at the end of this section. The input current from each presynaptic neuron *j* to each postsynaptic neuron *k* is modelled as the product of a linear function of the postsynaptic membrane potential (i.e., *v − x*_*k*_(*t*)) and the following nonlinear (sigmoid) function of the presynaptic membrane potential *x*_*j*_(*t*) [51]:

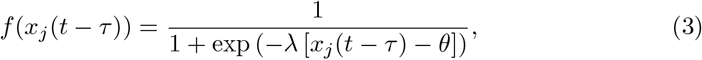

where *λ* is a positive constant that controls the shape of the sigmoid function and *θ* is the synaptic firing threshold [51, 65]. *τ* represents the transmission delay of the action potential along the chemical synapse (i.e., the time between depolarization of the presynaptic cell membrane and the input current received by the postsynaptic neuron). We set *λ* = 15, *θ* = −0.25, *τ* = 3.0 ms, and synaptic reversal potential *v* = 2.0 to model the excitatory coupling [51].

The topology of the neuronal network is defined by the chemical coupling connection matrix *C*, where element *c*_*jk*_ in Eq. (2a) is 1 if a synapse exists from neuron *j* to neuron *k* and is 0 otherwise. Because chemical synapses are unidirectional [62] and we assume no autapses (self-innervation), matrix *C* is asymmetric in general with zeros on the main diagonal. In this study, *N* = 100 neurons are arranged along a line of length *L* = 5 mm. This length was selected to represent the short axis of an ellipsoidal approximation of the STN based on experimental measurements of STN tissue volume [39, 46, 61]. The positions of the neurons are represented as points along line *L*, which are selected randomly from *N* intervals of equal length while ensuring a gap of at least 0.02 mm between points (the approximate diameter of the soma) for biological plausibility [39, 66]. We assume that the probability of a synaptic connection existing between neurons *j* and *k* (*P*_*jk*_) decreases with increasing inter-neuronal distance (*d*_*jk*_) [39]:

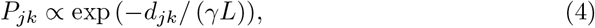

where *γ* is the probability decay constant, which determines whether synaptic connections are predominantly local (*γ* ≪ 1) or more homogeneous across the network (*γ* ≫ 1) [46]. In this work, we use *γ* = 0.24 for two reasons: (1) the resulting network has predominantly local connectivity, similar to the architecture used in existing studies [25, 40]; and (2) the average synaptic length is approximately 0.543 mm, as has been observed experimentally [39, 66]. Based on observations of the neuronal connectivity within the STN and following the methods of existing studies [39, 46, 67], 700 entries of the chemical coupling connection matrix *C* are set to 1 (i.e., 7% of all possible connections including autapses are activated in the network), all of which have probability *P*_*jk*_ ≥ 0.33 [39]. The connections are selected using a strategy analogous to simple fitness-proportionate (roulette-wheel) selection in evolutionary algorithms [68]. The selection process begins by setting all elements of *C* to 0, computing *P*_*jk*_ for *j* ≠ *k*, and zeroing all computed probabilities *P*_*jk*_ *<* 0.33. A cumulative distribution function is then constructed and a random number is drawn uniformly to activate one of the possible connections according to their relative probabilities. The probability associated with the selected connection is then zeroed and the process is repeated until 700 connections have been selected.

The strengths of the synapses (weights *w*_*jk*_(*t*) in Eq. (2a)) are updated following the STDP rule. The connection weight *w*_*jk*_(*t*) represents the strength of the chemical synapse from presynaptic neuron *j* to postsynaptic neuron *k*; we use the variables *t*_pre_ and *t*_post_ to represent their most recent spike times. Spikes are identified from the action potentials *x*(*t*) using two criteria. First, the neuron’s action potential must cross a threshold [51]; in this study, we use a threshold of 1.5, which was determined based on the values of the (dimensionless) potentials *x*(*t*) in our network. Second, after having crossed the threshold, the action potential must drop to near the resting potential (*x*_0_ + *ϵ*, where *ϵ* = 0.1) before another spike can be registered. Therefore, in the case of bursting behaviour, only the first spike within the burst is detected, which is consistent with previous studies of abnormal rhythmic bursting activity in Parkinson’s disease [58, 69, 70]. Immediately after a spike is detected in the presynaptic neuron, the postsynaptic neuron, or both, synaptic weight *w*_*jk*_(*t*) is incremented by Δ*w* as follows [25]:

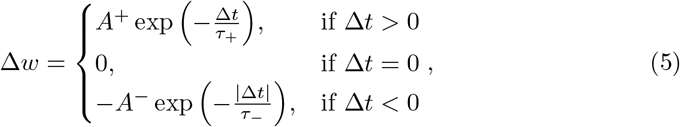

where Δ*t* = *t*_post_ − *t*_pre_ − *t*_d_, with synaptic transmission delay *t*_d_ set to 3.0 ms. Parameters *τ*_+_ and *τ*− represent, respectively, the decay time constants for long-term potentiation (Δ*w >* 0) and long-term depression (Δ*w <* 0) in the STDP rule. We set *A*^+^ = 0.00003 to implement a slow STDP rule that ensures smooth synaptic weight updates in our network of 100 neurons, *τ*_+_ = 10 ms, and *τ*− = 40 ms, adopting values used in the literature [25, 52]. Finally, we set *A*^*−*^ = *A*^+^*β/τ*_R_, where *β* = 1.9 is the ratio of overall depression to potentiation (when *β >* 1, long-term depression dominates over long-term potentiation [40]) and *τ*_R_ = *τ*− */τ*_+_ [25]. In our simulations, we clamp the synaptic weights to impose the bounds *w*_*jk*_(*t*) ∈ [0, 1].

Parameter *σ* in Eq. (2a) scales the influence of presynaptic neuronal activity on postsynaptic neurons. In our network of 100 neurons with 7% connectivity, there are 700 synapses in total and, therefore, 7 incoming synapses per neuron on average (represented by the braced expression in Eq. (2a)). The braced expression in Eq. (2a) may be approximated assuming (*v* − *x*_*k*_(*t*)) ∈ [0.5, 3.5] since *v* = 2.0 and *x*_*k*_(*t*) ∈ [−1.5, 1.5], weights *w*_*jk*_(*t*) ≈ 0.4, *c*_*jk*_ ∈ {0, 1} (on average, 7 of which are 1 for each neuron *k*), and noting that *f* (*x*_*j*_(*t* − *τ*)) ∈ [0, 1] for each incoming synapse. Under these assumptions, the value of the braced expression in Eq. (2a) is maximally 9.8*σ*. Large values of *σ* would cause the braced expression to exceed the physiologically plausible range of input associated with spiking behaviour in the STN. We tuned *σ* to prevent excessive synaptic inputs, setting *σ* = 0.2 in this study. The values we used for the parameters in the HR model and STDP rule are summarized in Tables 1 and 2, respectively.

**Table 1.** Parameter values used in the Hindmarsh–Rose (HR) neuronal model.

| Parameter | Value | Description |
| --- | --- | --- |
| $b$ | 3.0 | Regulates neuron’s dynamics (spiking or bursting) |
| $i^{\text{ext}}$ | 1.45 | External current injected into the neuron |
| $L$ | 5 mm | Length of neuronal network |
| $N$ | 100 | Number of neurons |
| $r$ | 0.01 | Regulates the spiking frequency |
| $v$ | 2.0 | Synaptic reversal potential |
| $x_0$ | −1.6 | Resting potential |
| $\gamma$ | 0.24 | Probability decay constant for determining synaptic connections |
| $\theta$ | −0.25 | Synaptic firing threshold |
| $\lambda$ | 15 | Controls shape of sigmoid function in fast-threshold-modulation model |
| $\sigma$ | 0.2 | Synaptic input scaling factor |
| $\tau$ | 3.0 ms | Synaptic transmission delay |

**Table 2.** Parameter values used in the spike-timing-dependent plasticity (STDP) rule.

| Parameter | Value | Description |
| --- | --- | --- |
| $A^+$ | 0.00003 | Potential update amplitude |
| $A^-$ | $A^+ \beta / \tau_R$ | Depression update amplitude |
| $t_d$ | 3.0 ms | Synaptic transmission delay |
| $\beta$ | 1.9 | Depression-to-potential ratio |
| $\tau_+$ | 10 ms | Decay time constant for long-term potentiation |
| $\tau_-$ | 40 ms | Decay time constant for long-term depression |
| $\tau_R$ | $\tau_- / \tau_+$ | Ratio of decay time constants |

### 2.2 Order parameter

The Kuramoto order parameter quantifies the degree of spike synchrony in a neuronal network [71]. To compute the Kuramoto order parameter at each simulation time point *t*, the phase of each neuron (*ϕ*_*k*_(*t*)) is approximated as follows [72]:

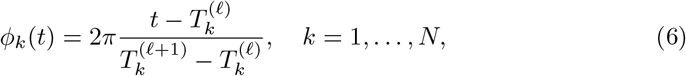

where 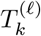 and 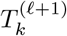 are, respectively, the time points at which spikes *l* and *l* + 1 were detected for neuron *k*, and 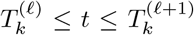. The Kuramoto order parameter (*ρ*(*t*)) reflects the average coherence among all neuronal phases [51]:

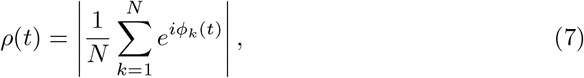

Where 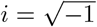. For visual clarity, we compute a moving average of Kuramoto order parameter 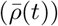 over 10-second intervals:

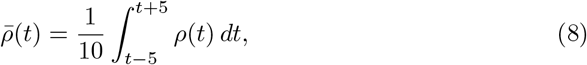

where 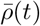 is sampled every 0.1 s. Low values of 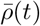 (near 0) indicate weak phase coherence while high values (near 1) indicate strong coherence and a synchronized network.

### 2.3 Coordinated reset stimulation

To study the effect of RVS CR on the network’s neuronal spike synchrony, a stimulation current 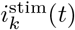 is added to Eq. (2a):

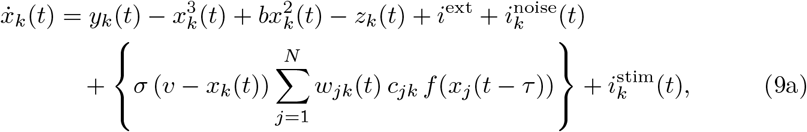

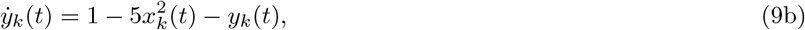

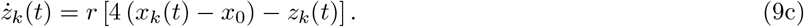

The RVS CR stimulation protocol has been described previously [25, 40, 73, 74]. Electrical stimulation is delivered to the tissue as a sequence of pulses of specified amplitude (pulse height) and duration (pulse width) [40, 75]. In this study, we tuned the stimulation parameters such that the pulses were of sufficient amplitude and sufficiently brief to reliably evoke a spike response [25]. In clinical DBS treatment, electrical stimuli are typically charge balanced to prevent tissue damage from charge accumulation and electromechanical reactions at the electrodes [76]. Charge balancing involves delivering pulses in pairs: an excitatory pulse (a high-amplitude current that induces neuronal spiking) followed by an inhibitory pulse (a low-amplitude current of opposite polarity that draws accumulated positive ions back toward the electrode) [25, 40, 76]. The low amplitude of the inhibitory pulse is unlikely to substantially influence neuronal spiking behaviour. Provided the stimulation is sufficiently strong and fast, spikes occur primarily in response to stimulation [25] and stimulus-induced spikes dominate the phase resetting mechanism. Thus, for simplicity and following the methods of Liu et al. [48], we apply only excitatory pulses in our simulations.

In the RVS CR technique, stimulation is applied sequentially to distinct subpopulations of neurons (stimulation sites) surrounding separate electrode contacts. This technique differs from conventional local square-wave perturbation approaches, where the stimulation is typically applied repeatedly to the same neuronal population to perturb local neuronal activity [77, 78]. Following the methods of Kromer et al. [25], we define subpopulation *m* as comprising neurons in the interval [*L* (*m* − 1) */S, Lm/S*], *m* = 1, …, *S*, where *L* = 5 mm and *S* is the number of subpopulations. In each RVS CR stimulation cycle, each subpopulation receives exactly one excitatory pulse. Pulses are delivered to one subpopulation at a time in a random sequence. The period of each RVS CR stimulation cycle is 1*/f*_CR_, where *f*_CR_ is the stimulation (cycle) frequency; therefore, an excitatory pulse begins every 1*/* (*Sf*_CR_) seconds [25, 40] (Fig. 1).

**Fig. 1.**
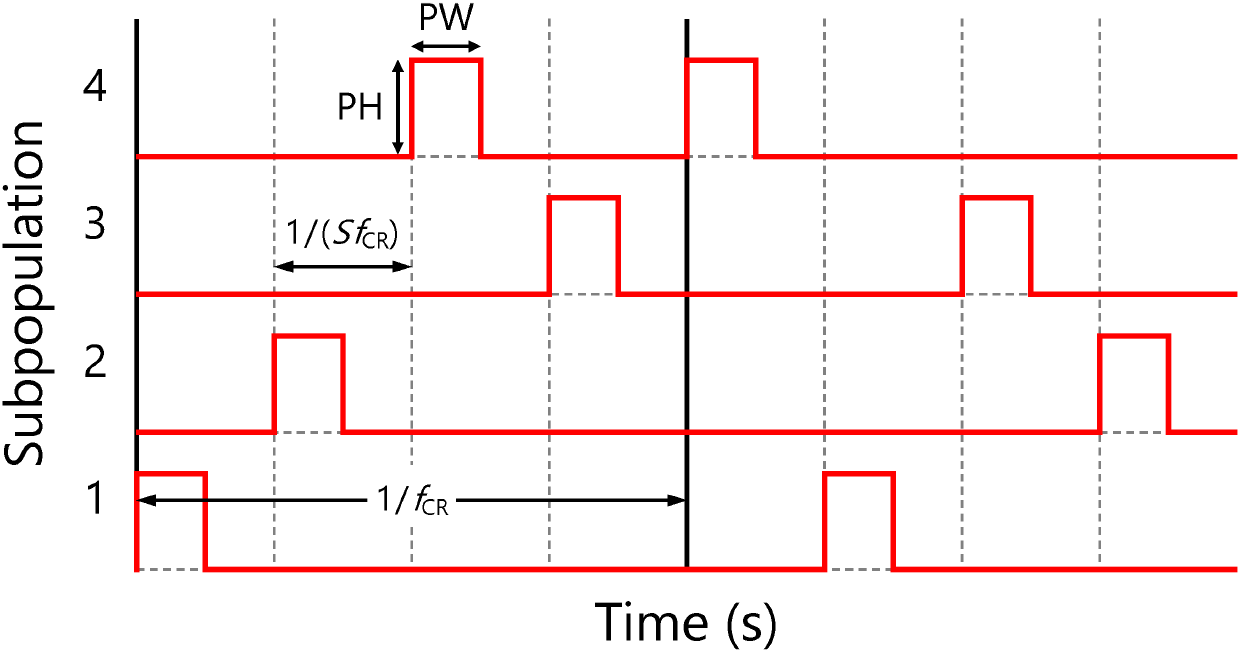
Example of coordinated reset stimulation with rapidly varying sequences (RVS CR) with *S* = 4 subpopulations of neurons. PH: pulse height, PW: pulse width, *f*_CR_: stimulation frequency.

### 2.4 Numerical experiments

Simulations are performed in MATLAB R2020b and implemented as a MEX function using the Microsoft Visual C++ 2019 compiler for efficiency. The fourth-order Runge–Kutta (RK4) method is used to solve Eq. (9) with *N* = 100 neurons, using a fixed integration step size of 0.1 ms. Simulations of duration 200 s are performed to compare the effects of uniformly distributed white noise and Poisson noise on the system’s mean synaptic weight and order parameter. To test the robustness of our observations, we repeat these simulations using low, moderate, and high initial conditions for the synaptic weights and using five random network topologies. Each of these five networks is generated using the same connection probability but with different random seeds, resulting in different connectivity patterns (i.e., different *C* matrices). The evolution of individual synaptic weights is analyzed for the low- and high-initial-weight conditions using the network with white noise to verify our system’s behaviour. To explore the sensitivity of RVS CR stimulation to the number of stimulation sites, we generate simulations of duration 1400 s with 2, 12, and 24 subpopulations, and compare the resulting mean synaptic weights and order parameters over time. Finally, we validate our results against the simulations of Kromer et al. [25] over a large range of subpopulations and stimulation frequencies. For this comparison, initial synaptic weights are selected randomly from a uniform distribution over the range [0.45, 0.55] so that the network’s mean synaptic weight quickly stabilizes. Simulations of duration 2000 s are generated, with RVS CR stimulation starting at 200 s. Pulse height 20 and pulse width 0.6 ms ensure that spikes are reliably induced while retaining sufficient temporal separation even at high stimulation frequencies. At lower amplitudes, the effect of the stimulation resembles that of noise, where the applied pulses primarily shift the timing of the intrinsic neuronal spiking rather than inducing phase resetting. One simulation is generated for each combination of subpopulation number and stimulation frequency, and the mean synaptic weight is computed over the final 100 s of each simulation (to average out persisting fluctuations after the main changes in the mean synaptic weight have occurred).

## 3 Results and discussion

We adopt the HR neuronal network model with unidirectional excitatory chemical synapses, with the evolution of the synaptic weights governed by the STDP rule. In this section, we compare the effect of two types of noise in the differential equation representing the neuronal membrane potential, we examine the evolution of synaptic weights in detail, and we apply RVS CR stimulation to demonstrate the utility of the proposed model for studying this treatment strategy.

### 3.1 Effect of noise model on network behaviour

Uniformly distributed white noise and Poisson noise are both used in the existing literature. As described above, we generated simulations of 200 s duration using five random network topologies, each comprising 100 neurons and 700 synapses. The evolution of the mean synaptic weights and order parameters are shown in Fig. 2 when using uniformly distributed white noise in Eq. (2a). As shown, three simulations were generated using each network topology, where all initial synaptic weights were either set to zero or selected randomly from a uniform distribution over the range [0.35, 0.45] or [0.75, 0.85]. Regardless of the initial synaptic weights, the mean weight converged to approximately 0.4 and the order parameter converged to approximately 0.8. Although the order parameter converged to a value less than 1, the network is indeed in a synchronized state, as evidenced by coherent spike timing in the raster plot (see Fig. 3, below). Unlike LIF models, the HR model exhibits bursting behaviour, chaotic dynamics, and stochastic fluctuations that prevent perfect phase alignment. Therefore, the order parameter is less than 1 even when the network is in a stable synchronized state. In all cases, neuronal synchrony emerged following an initial transient phase. The noise caused persistent fluctuations in both the mean synaptic weight and the order parameter, but these fluctuations were of insufficient magnitude to substantially shift the spike timing or disrupt the system’s inherent tendency to synchronize. Raster plots showing the neuronal spiking behaviour in one randomly generated network are shown in Fig. 3 for each of the three initial synaptic weight conditions. With initial synaptic weights of zero, the neurons were initially desynchronized (Fig. 3(a)) due to their randomly initialized states and the influence of the white noise on the spiking behaviour at low synaptic weights; the STDP rule increased the mean synaptic weight over time, driving the system to synchrony (Fig. 3(b)). With moderate or high initial synaptic weights (Figs. 3(c) and 3(e)), the neurons were initially synchronized, though at different dominant frequencies (i.e., firing at different rates). The dominant frequency was lower (approximately 5 Hz) when the initial synaptic weights were higher (Fig. 3(e)) because postsynaptic neurons were driven into a bursting regime by the greater synaptic inputs; our algorithm for detecting spikes registered only the first spike in each burst. Regardless of the initial synaptic weights, the dominant frequency of our network was approximately 8.5 Hz once it had reached its stable synchronized state. Previous studies have reported that tremor symptoms in PD are associated with pronounced synchronization in the STN around the theta–alpha frequency band (4–10 Hz) [25, 79, 80]; thus, our model is appropriately tuned for studying tremor-related treatments in PD.

**Fig. 2.**
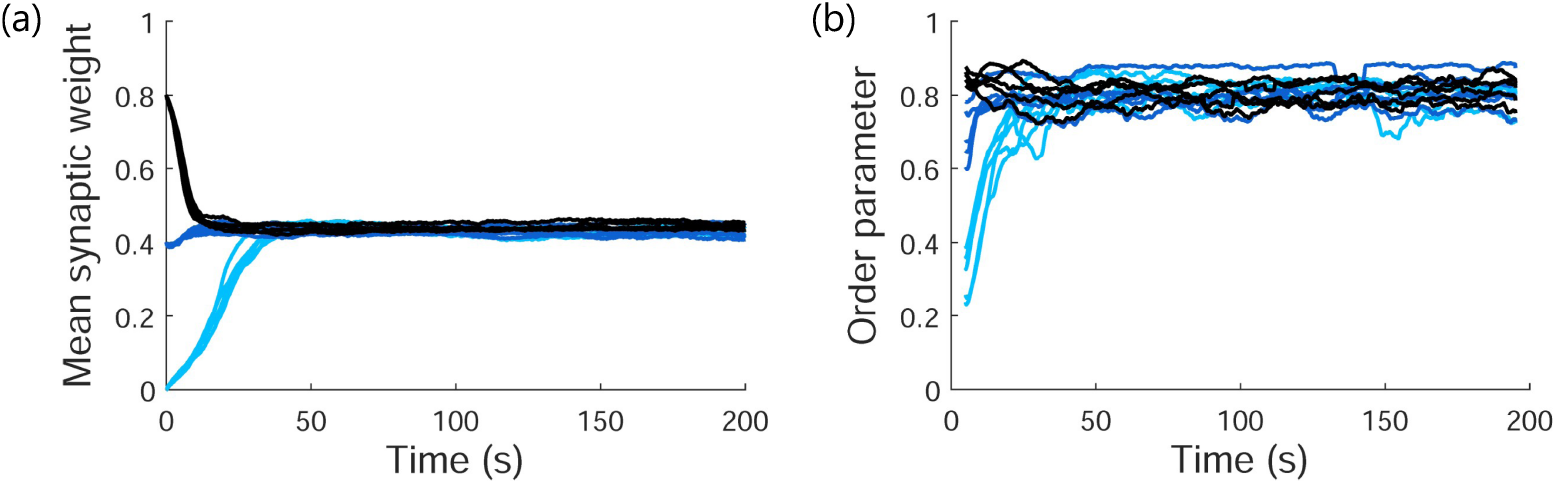
Network behaviour with uniformly distributed white noise. Evolution of (a) mean synaptic weight and (b) time-averaged Kuramoto order parameter 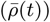 in five random networks, each simulated with initial synaptic weights equal to zero (light blue), randomly selected between 0.35 and 0.45 (dark blue), and randomly selected between 0.75 and 0.85 (black). All networks exhibited inherent neural synchrony.

**Fig. 3.**
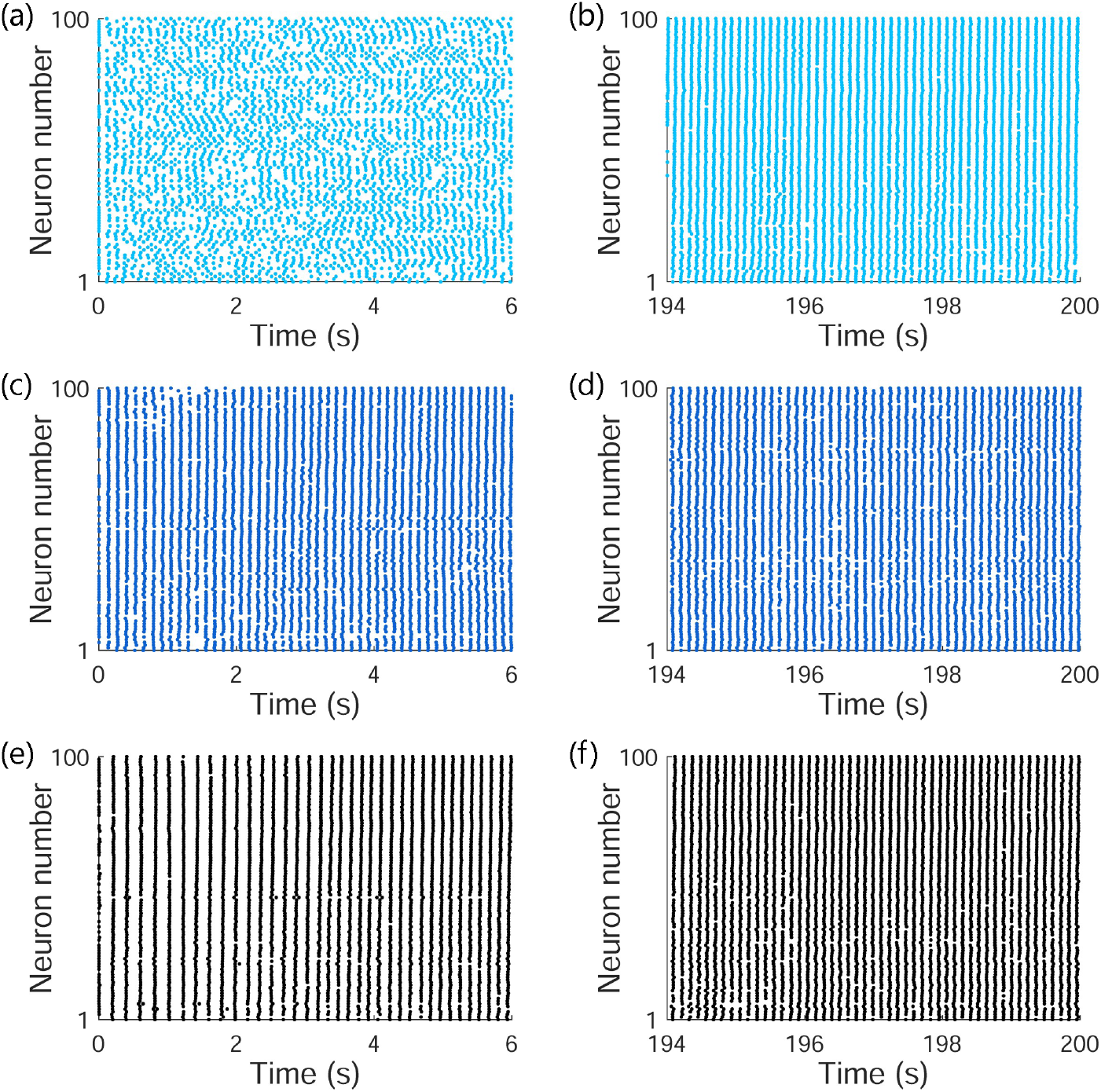
Raster plots of neuronal spiking activity in one randomly generated network. First and last 6 s of 200 s simulations with different initial synaptic weights: zero ((a) and (b)), randomly selected between 0.35 and 0.45 ((c) and (d)), and randomly selected between 0.75 and 0.85 ((e) and (f)). In all cases, the system stabilizes in a synchronized state with a dominant frequency of approximately 8.5 Hz. Note that in the case of bursting behaviour, only the first spike within a burst is shown.

The evolution of the mean synaptic weights and order parameters are shown in Fig. 4 when using Poisson noise in Eq. (2a). Four simulations were generated using each of five random network topologies, where all initial synaptic weights were selected randomly from a uniform distribution over the range [0, 0.1], [0.15, 0.25], [0.45, 0.55], or [0.75, 0.85]. As shown, the mean synaptic weight converged close to 0 (a stable, desynchronized state) when initial weights were low and to approximately 0.5 (a stable, synchronized state) when the initial weights were high. In contrast to uniformly distributed white noise, Poisson noise induces shifts in the spike timing of the HR model that disrupt the intrinsic rhythm of the network when the mean synaptic weight is low. Consequently, when initial synaptic weights were low, the mean synaptic weight remained low and the network remained desynchronized. The existence of two stable states (one desynchronized and one synchronized) when using Poisson noise may be useful in some analytical contexts; however, in this study, the noise model has little effect on the network’s behaviour during the application of CR stimulation because the stimulation amplitude is sufficiently high to reliably evoke a spike in response to each stimulation pulse. Consequently, neuronal firing is predominantly determined by the CR stimulation pattern. The evolution of the mean synaptic weight during stimulation, which is the focus of the present study, is only weakly affected by the details of the noise model, as corroborated by our preliminary analyses. Furthermore, Poisson noise requires an additional differential equation and is, therefore, more computationally expensive than a white noise model. Therefore, in the remainder of this study, we proceed with uniformly distributed white noise and a network with a single stable synchronized state, and we do not consider the network’s behaviour following cessation of CR stimulation, where the existence or nonexistence of bistability would be relevant. Nevertheless, in studies that consider the post-stimulation dynamics, employing the Poisson noise model may increase physiological accuracy.

**Fig. 4.**
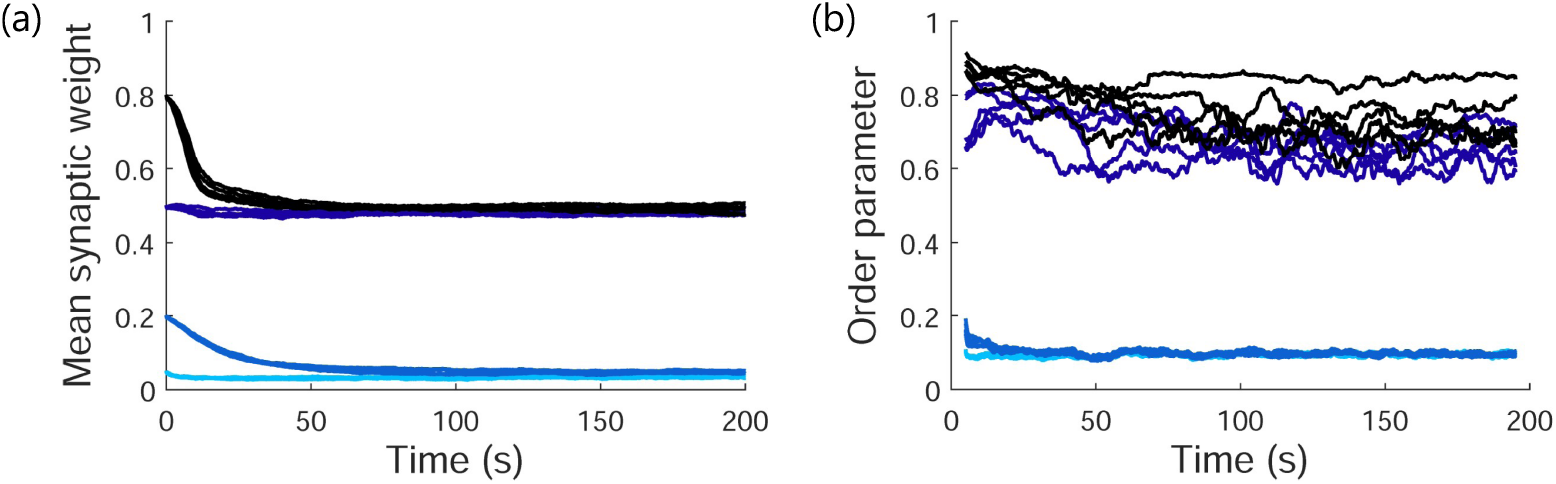
Network behaviour with Poisson noise. Evolution of (a) mean synaptic weight and (b) time-averaged Kuramoto order parameter 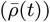 in five random networks, each simulated with initial synaptic weights that were randomly selected between 0 and 0.1 (light blue), 0.15 and 0.25 (blue), 0.45 and 0.55 (dark blue), or 0.75 and 0.85 (black). Each network stabilized in either a desynchronized state (low mean synaptic weight) or a synchronized state (high mean synaptic weight).

### 3.2 Dynamics of synaptic weights

The proposed HR model with STDP permits detailed investigation of neuronal network dynamics. As an example, we consider a network comprising 100 neurons and 700 synapses with random topology and uniformly distributed white noise. Two simulations of 200 s duration were generated: one with low initial synaptic weights (selected randomly between 0.01 and 0.05) and one with high initial synaptic weights (between 0.75 and 0.85). The spiking activity of one neuron in the middle of the network and all of its presynaptic neurons is shown in Fig. 5; the dynamics of the corresponding synaptic weights are illustrated in Fig. 6. As described in Section 2.1, the STDP rule updates synaptic weights based on the relative spike timing of the presynaptic and postsynaptic neurons. Specifically, if the presynaptic neuron spikes *before* the postsynaptic neuron, the synaptic weight *increases* (long-term potentiation); if the presynaptic neuron spikes *after* the postsynaptic neuron, the synaptic weight *decreases* (long-term depression) [38, 81]. These dynamics are evident in Figs. 5 and 6. For example, in Fig. 5(b), presynaptic neuron 73 was spiking after postsynaptic neuron 50 during the last second of the simulation with low initial synaptic weights; consequently, in Fig. 6(f), we observe that the weight of the synapse from neuron 73 to neuron 50 was low at the end of this simulation (Fig. 6(f), light blue line). The opposite was true in the simulation with high initial synaptic weights: presynaptic neuron 73 was spiking before postsynaptic neuron 50 during the last second of the simulation (Fig. 5(d)) so the corresponding synaptic weight was high at the end of the simulation (Fig. 6(f), black line). In contrast, presynaptic neuron 24 was spiking before postsynaptic neuron 50 in Fig. 5(b) but after neuron 50 in Fig. 5(d); consequently, in Fig. 6(a), we observe that the corresponding synaptic weight was high at the end of the low-initial-weight simulation but low at the end of the high-initial-weight simulation. An important observation is that, although the mean synaptic weight and time-averaged Kuramoto order parameter have settled after 200 s (see Fig. 2), variations in some of the individual synaptic weights continued due to the STDP rule and the noise term (*i*^noise^(*t*)) in Eq. (2a).

**Fig. 5.**
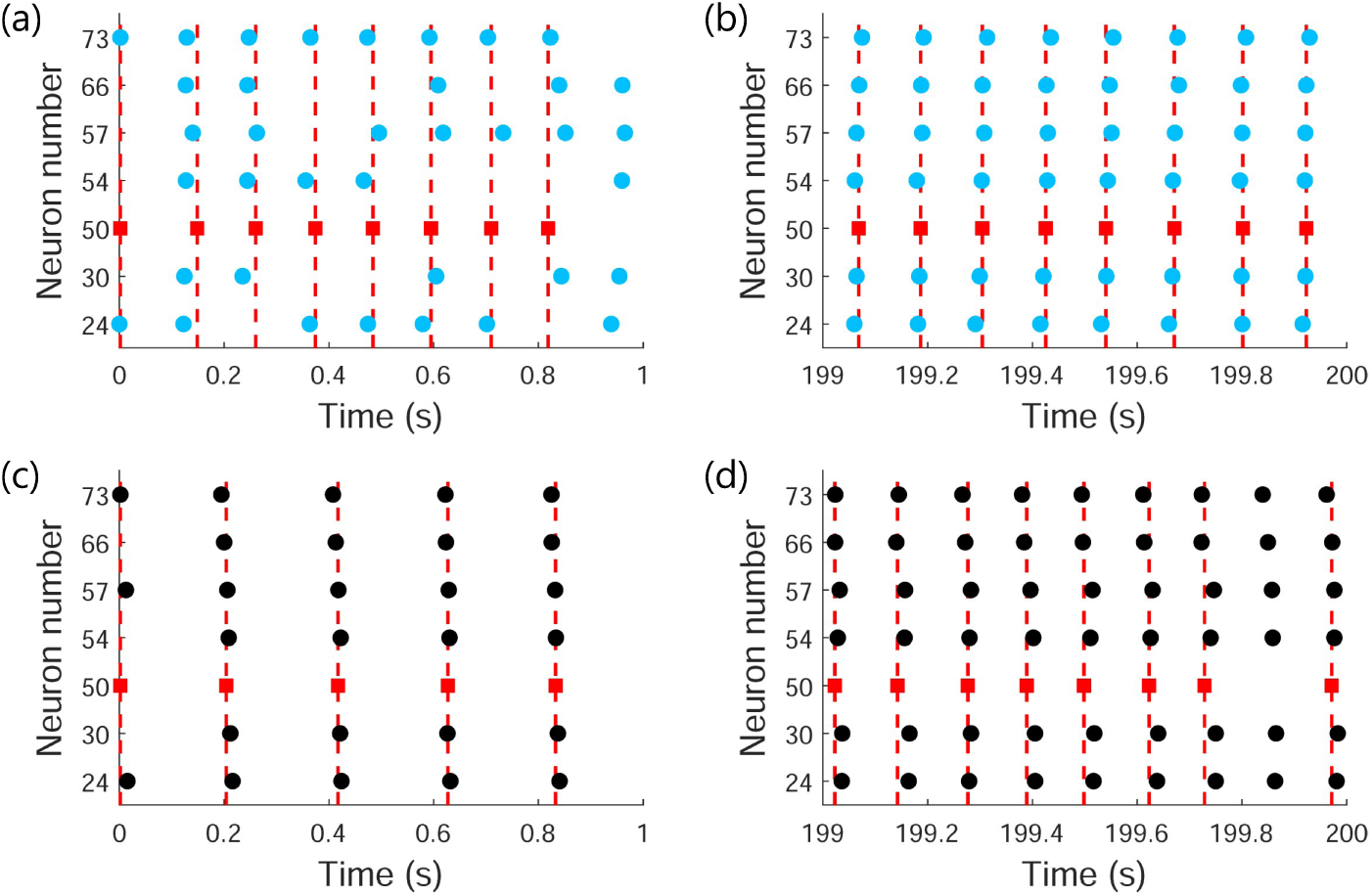
Raster plots of spiking activity for one postsynaptic neuron (number 50; red squares) and all of its presynaptic neurons (24, 30, 54, 57, 66, and 73; circles) in one randomly generated network. Spiking behaviour is shown from a simulation with low initial synaptic weights (light blue) during the (a) first second and (b) last second of the simulation, and from a simulation with high initial synaptic weights (black) during the (c) first second and (d) last second of the simulation. Dashed vertical lines aligning with the spikes of the postsynaptic neuron (number 50) are shown for reference.

**Fig. 6.**
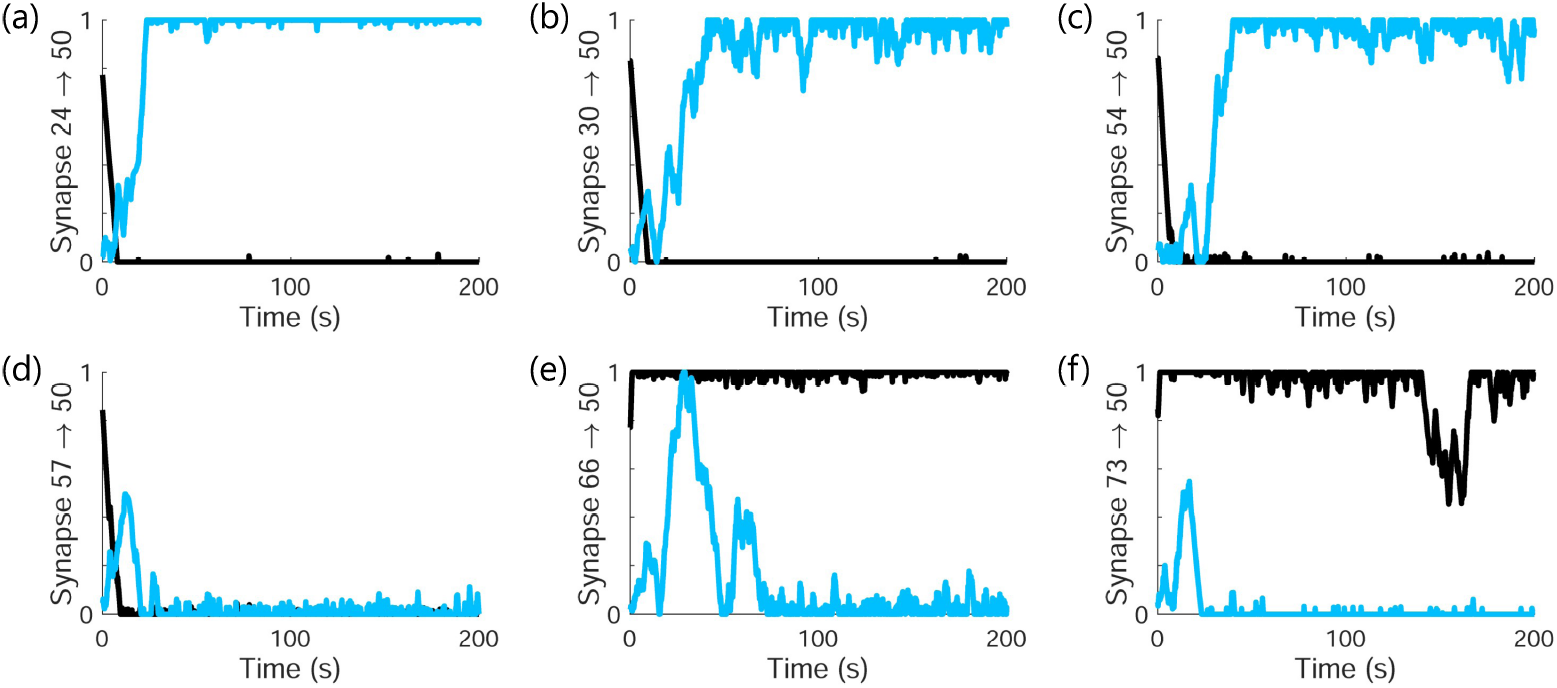
Evolution of synaptic weights corresponding to the raster plots shown in Fig. 5. Six synapses project onto postsynaptic neuron 50 in the randomly generated network; panels (a)–(f) illustrate the corresponding synaptic weights during a simulation with low initial synaptic weights (light blue) and a simulation with high initial synaptic weights (black).

### 3.3 Model verification

RVS CR stimulation was applied to the proposed HR model with STDP, as described in Eq. (9). We consider a network comprising 100 neurons and 700 synapses with random topology and uniformly distributed white noise, with a dominant frequency of 8.5 Hz. As an example, Fig. 7 illustrates the mean synaptic weight and order parameter for simulations of duration 1400 s, where CR stimulation was configured with pulse height 20, pulse width 0.6 ms, and frequency 23 Hz; the corresponding raster plots from 2 s before to 2 s after the onset of stimulation are shown in Fig. 8. The initial synaptic weights were selected randomly between 0.45 and 0.55; CR stimulation commenced after 200 s to allow the initial transients to decay. As shown, the mean synaptic weight and order parameter decreased dramatically when CR stimulation was applied and the neuronal population was divided into 2 or 24 subpopulations (stimulation sites), indicating a reduction in the synchronization of the network and success of the RVS CR therapy. The mean synaptic weight decreased the fastest with 2 subpopulations. However, when the neuronal population was divided into 12 subpopulations, the mean synaptic weight increased following a brief decrease at the onset of the stimulation.

**Fig. 7.**
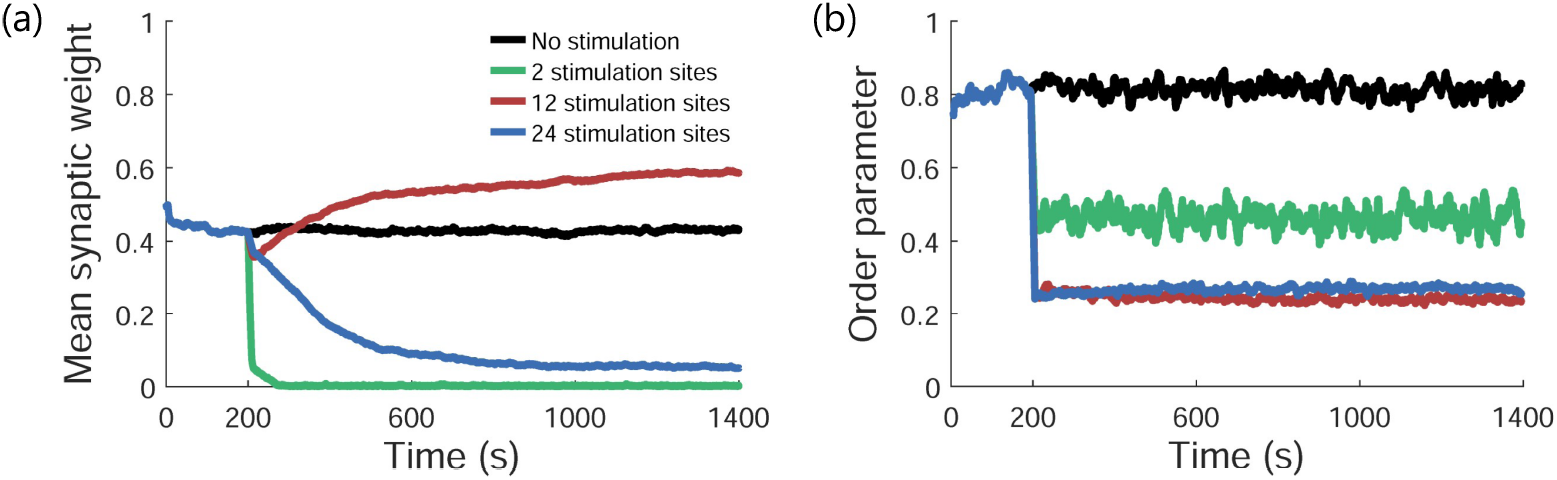
The effectiveness of coordinated reset stimulation at decreasing neuronal synchrony depends on the number of stimulation sites (i.e., the number of neuronal subpopulations that receive stimulation simultaneously). Decreases in (a) mean synaptic weight and (b) order parameter are observed with 2 and 24 stimulation sites following onset of stimulation at 200 s, indicating successful desynchronization, but mean synaptic weight increases with 12 stimulation sites. Results are shown for a stimulation frequency of 23 Hz.

**Fig. 8.**
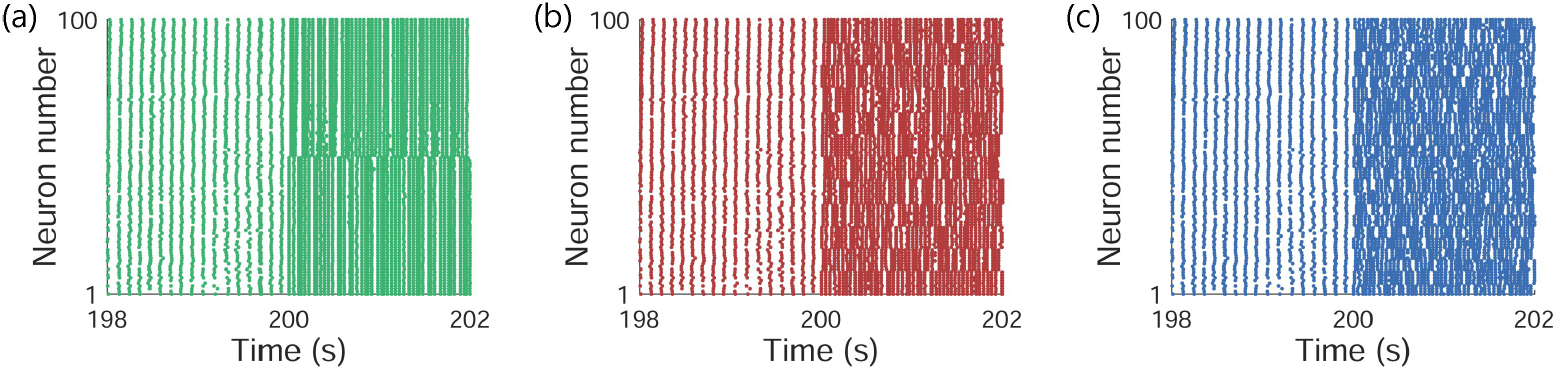
Raster plots of neuronal spiking activity from 2 s before to 2 s after onset of coordinated reset stimulation. The network is initially synchronized; stimulation causes each subpopulation of neurons to spike together. Stimulation is applied to (a) 2 sites, (b) 12 sites, or (c) 24 sites.

Although the order parameter still decreased in this case, the increase in the mean synaptic weight indicates that neural synchrony increased, suggesting that the RVS CR therapy would be ineffective with this configuration. A similar result was reported by Kromer et al. [25], who used an LIF neuronal model comprising 1000 neurons and 70,000 synapses (7% of all possible connections, consistent with the connectivity observed in the STN [67] as well as the connectivity of our network), with a dominant frequency of 3.5 Hz. Specifically, they observed a decrease in the mean synaptic weight with 2 and 24 subpopulations but an increase with 12 subpopulations. We note that this result was observed in their model when using a different stimulation frequency (17.5 Hz), which may be attributed to differences in the network size (number of neurons and synapses) and, consequently, the ratio of depression to potentiation (*β*). We discuss these differences in more detail below. Nevertheless, our proposed model demonstrates the same behaviour with only 10% as many neurons and 1% as many synapses.

To further validate our proposed model, we explored the response of the network to RVS CR stimulation with between 2 and 22 stimulation sites (*S*) and using stimulation frequency (*f*_CR_) between 2 and 50 Hz. In total, 550 simulations were generated, each of duration 2000 s with RVS CR stimulation starting at 200 s. As shown in Fig. 9(a), the model predicted poor efficacy of RVS CR therapy for certain combinations of these parameters. For example, if 6 stimulation sites were used, CR stimulation would effectively reduce the mean synaptic weight with low stimulation frequencies (blue squares in Fig. 9(a)) but would be ineffective at desynchronizing the network with stimulation frequencies above approximately 24 Hz. The dashed curves in Fig. 9(a) are defined by 1*/* (*Sf*_CR_) = *t*_d_*/k* for *k* ∈ {1, 2, 3} and *t*_d_ = 0.003 s. The lower of the three dashed curves in Fig. 9(a) represents the boundary at which the spike-timing difference (*t*_post_ − *t*_pre_) between postsynaptic and presynaptic neurons of inter-subpopulation synapses (i.e., synapses between neurons belonging to distinct stimulation sites) is equal to *t*_d_. According to the STDP rule (Eq. (5)), spike-timing differences greater than *t*_d_ lead to synaptic potentiation. Consequently, CR stimulation leads to an increase in the network’s mean synaptic weight and is ineffective in regions of the parameter space that lie below this curve. The middle of the three dashed curves in Fig. 9(a) describes a similar phenomenon, but where the spike-timing difference is equal to *t*_d_*/*2. Here, the dynamics of the synaptic weights are governed not only by spike pairs within the same stimulation cycle, but also by spike pairs occurring within consecutive cycles [40]. In Fig. 9(b), we illustrate the effect of decreasing the depression-to-potentiation ratio (*β*) from 1.9 to 1.85. Changes in dopamine levels in the substantia nigra pars compacta can reshape the asymmetry of the STDP learning window [82], which can be modelled by changing the depression-to-potentiation ratio in our model. The shapes of the ineffective regions in this parameter space and their relationships with the dashed curves are consistent with the results of Kromer et al. [25], who used a much larger LIF neuronal model and set *β* = 1.4 to stabilize their network with a mean synaptic weight of approximately 0.4. In Fig. 9(c), we illustrate the evolution of the mean synaptic weight with 16 stimulation sites and using stimulation frequencies of 6 Hz, 26 Hz, and 46 Hz as an example. Although all three stimulation frequencies elicit a reduction in the mean synaptic weight, higher-frequency stimulation desynchronizes the network faster due to more frequent stimulation-induced spikes. In general, higher-frequency stimulation produces faster synaptic weight adaptation through the STDP mechanism. The results of Fig. 9 highlight the importance of tuning *β* when reconciling simulations with experimental data: *β* may vary across patients or over time and plays a key role in determining the predicted efficacy of CR stimulation.

**Fig. 9.**
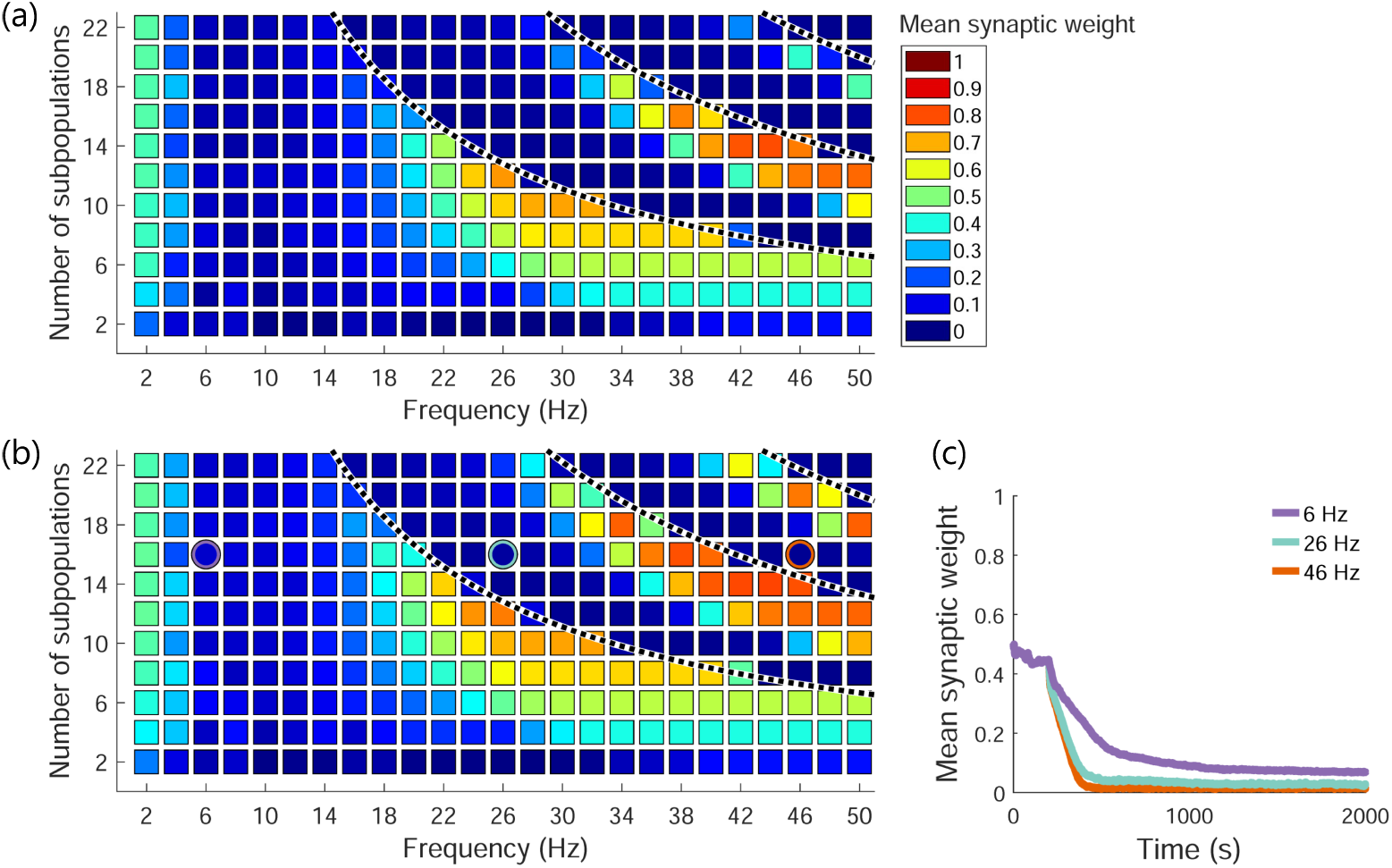
Effectiveness of coordinated reset stimulation as the number of stimulation sites and the stimulation frequency vary. Mean synaptic weight averaged over the last 100 s of a 2000 s simulation for a network with depression-to-potentiation ratio of (a) *β* = 1.9 and (b) *β* = 1.85, and (c) evolution of mean synaptic weight for the three simulations highlighted with circles in panel (b). The dashed curves in panels (a) and (b) represent the boundaries at which the spike-timing difference between postsynaptic and presynaptic neurons of inter-subpopulation synapses is equal to *t*_d_*/k* for *k* ∈ {1, 2, 3}.

## 4 Conclusion

In this study, we generated simulations of neuronal networks comprising 100 Hindmarsh–Rose (HR) neurons with unidirectional excitatory chemical synapses, where synaptic weights were updated according to a spike-timing-dependent plasticity (STDP) rule. Network topologies were generated randomly with 700 synapses (7% of all possible connections) based on observations of the neuronal connectivity within the subthalamic nucleus (STN). Numerical studies were performed to determine whether the proposed model is a viable alternative to the leaky integrate-and-fire neuronal models that appear in the existing literature, which cannot reproduce realistic spiking patterns [44, 50]. We demonstrated that including white noise in the HR model gives rise to a single stable state in which neurons are synchronized, with a dominant frequency of approximately 8.5 Hz. Regardless of the initial synaptic weights (whether initialized to low or high values), the mean synaptic weight converges to approximately 0.4, though the weight of each synapse remains dynamic due to the STDP rule and the white noise representing the spiking behaviour of unmodelled STN neurons. If Poisson noise is included in the HR model instead, the network will evolve toward one of two states: if initial synaptic weights are low, the mean synaptic weight will remain low and neurons will be desynchronized; alternatively, if initial synaptic weights are high, the mean synaptic weight will remain high and neurons will be synchronized. Finally, we applied the proposed model to study the efficacy of coordinated reset (CR) stimulation with rapidly varying sequences at desynchronizing a synchronized network.

This study has several important limitations. First, the network comprised 100 STN neurons arranged along a line. Our results corroborate those reported previously [25]; however, we did not determine the minimum network size required to produce the observed behaviour, we did not explore other arrangements of the neurons (e.g., within a square [83]), and we did not explicitly model neurons in other brain regions (e.g., the globus pallidus externus [39]). Second, the model includes many numerical parameters that are likely to vary among individuals and even among neurons within the STN. A sensitivity analysis with respect to these parameters may facilitate clinical translation, as the most robust computational results may indicate treatments that would be most widely effective. Third, all subpopulations in the network received CR stimulation and only excitatory pulses were applied. Again to facilitate clinical translation, it would be valuable to explore the effects of CR stimulation applied to only a subset of the STN neurons and to study the effect of inhibitory pulses subsequent to the excitatory pulses.

To the best of our knowledge, this is the first study to use an HR model with STDP for examining the effects of noise and the efficacy of CR stimulation. We tuned our model to represent synchronized neuronal activity associated with parkinsonian tremor, then generated simulations over a range of CR stimulation sites and frequencies, observing a complex pattern of effective and ineffective regions in this parameter space. We also demonstrated the sensitivity of this model to the depression-o-potentiation ratio (*β*) parameter in the STDP rule. Our results corroborate those of Kromer et al. [25], who used a large leaky integrate-and-fire neuronal model with a different depression-to-potentiation ratio and a different dominant frequency. The proposed HR-with-STDP model predicts network behaviour that compares favourably with the mean synaptic weight and order parameter results reported elsewhere in the literature, but offers two substantial benefits. First, the proposed model comprises only 100 neurons and 700 synapses (compared to 1000 neurons and 70,000 synapses used by Kromer et al. [25]), enabling efficient systematic investigation of stimulation parameters while retaining nonlinear neuronal dynamics. Smaller models can facilitate detailed analyses while reducing computational burden, enabling researchers to efficiently explore a wide range of stimulation parameters. Second, the proposed model provides realistic spiking patterns, which can be exploited to enhance clinical interpretation and translation—for example, by enabling calculation of local field potentials. Future work includes performing a systematic sensitivity analysis with respect to the network size, model parameters, and stimulation pulse amplitude and duration; studying additional model outputs such as local field potentials and the statistical distribution of synaptic weights; applying the HR-with-STDP model to study personalization of CR stimulation strategies; simulating CR therapy for individuals with Parkinson’s disease presenting with different symptoms (e.g., tremor vs. bradykinesia and rigidity [84, 85]); and expanding the model to include other regions of the basal ganglia and thalamus [86].

## Acknowledgments

The authors thank Anthony Lee for providing feedback on preliminary results. This work was supported by the Government of Canada’s New Frontiers in Research Fund (NFRFE-2023-00258 [Thomas K. Uchida, Mazen Al Borno]) and the Natural Sciences and Engineering Research Council of Canada (RGPIN-2019-05726 [Thomas K. Uchida]). The funders had no role in study design, data collection and analysis, decision to publish, or preparation of the manuscript.

## Statements and Declarations

## Funding

This work was supported by the Government of Canada’s New Frontiers in Research Fund (NFRFE-2023-00258 [Thomas K. Uchida, Mazen Al Borno]) and the Natural Sciences and Engineering Research Council of Canada (RGPIN-2019-05726 [Thomas K. Uchida]). The funders had no role in study design, data collection and analysis, decision to publish, or preparation of the manuscript.

## Competing interests

The authors have declared that no competing interests exist.

## Author contributions

- **Shahin Sharafi:** Conceptualization, Methodology, Software, Formal analysis, Data curation, Writing – original draft, Writing – review and editing, Visualization.
- **Jesse I. Gilmer:** Conceptualization, Methodology, Writing – review and editing.
- **Mazen Al Borno:** Conceptualization, Methodology, Writing – review and editing, Supervision, Funding acquisition.
- **Thomas K. Uchida:** Conceptualization, Methodology, Software, Writing – review and editing, Visualization, Supervision, Funding acquisition.

All authors read and approved the final manuscript.

## Data availability

The manuscript contains all information necessary to reproduce the simulation results presented in this study.

## Notes

### Competing Interest Statement

The authors have declared no competing interest.

### Summary of Updates

The manuscript has been revised in response to feedback provided by reviewers. Specifically, two references were added to the Introduction and minor edits were made to the text in five places.

## References

[1] Dorsey, E.R., Elbaz, A., Nichols, E., Abbasi, N., Abd-Allah, F., Abdelalim, A., Adsuar, J.C., Ansha, M.G., Brayne, C., Choi, J.-Y.J., Collado-Mateo, D., Dahodwala, N., Do, H.P., Edessa, D., Endres, M., Fereshtehnejad, S.-M., Foreman, K.J., Gankpe, F.G., Gupta, R., Hamidi, S., Hankey, G.J., Hay, S.I., Hegazy, M.I., Hibstu, D.T., Kasaeian, A., Khader, Y., Khalil, I., Khang, Y.-H., Kim, Y.J., Kokubo, Y., Logroscino, G., Massano, J., Ibrahim, N.M., Mohammed, M.A., Mohammadi, A., Moradi-Lakeh, M., Naghavi, M., Nguyen, B.T., Nirayo, Y.L., Ogbo, F.A., Owolabi, M.O., Pereira, D.M., Postma, M.J., Qorbani, M., Rahman, M.A., Roba, K.T., Safari, H., Safiri, S., Satpathy, M., Sawhney, M., Shafieesabet, A., Shiferaw, M.S., Smith, M., Szoeke, C.E.I., Tabarés-Seisdedos, R., Truong, N.T., Ukwaja, K.N., Venketasubramanian, N., Villafaina, S., Weldegwergs, K.G., Westerman, R., Wijeratne, T., Winkler, A.S., Xuan, B.T., Yonemoto, N., Feigin, V.L., Vos, T., Murray, C.J.L.: Global, regional, and national burden of Parkinson’s disease, 1990–2016: a systematic analysis for the Global Burden of Disease Study 2016. Lancet Neurol. 17(11), 939–953 (2018). doi:10.1016/s1474-4422(18)30295-3

[2] Su, D., Cui, Y., He, C., Yin, P., Bai, R., Zhu, J., Lam, J.S.T., Zhang, J., Yan, R., Zheng, X., Wu, J., Zhao, D., Wang, A., Zhou, M., Feng, T.: Projections for prevalence of Parkinson’s disease and its driving factors in 195 countries and territories to 2050: modelling study of Global Burden of Disease Study 2021. BMJ 388, e080952 (2025). doi:10.1136/bmj-2024-080952

[3] Hirsch, E., Graybiel, A.M., Agid, Y.A.: Melanized dopaminergic neurons are differentially susceptible to degeneration in Parkinson’s disease. Nature 334(6180), 345–348 (1988). doi:10.1038/334345a0

[4] Höglinger, G.U., Rizk, P., Muriel, M.P., Duyckaerts, C., Oertel, W.H., Caille, I., Hirsch, E.C.: Dopamine depletion impairs precursor cell proliferation in Parkinson’s disease. Nat. Neurosci. 7(7), 726–735 (2004). doi:10.1038/nn1265

[5] Mitchell, I.J., Clarke, C.E., Boyce, S., Robertson, R.G., Peggs, D., Sambrook, M.A., Crossman, A.R.: Neural mechanisms underlying parkinsonian symptoms based upon regional uptake of 2-deoxyglucose in monkeys exposed to 1-methyl-4-phenyl-1,2,3,6-tetrahydropyridine. Neuroscience 32(1), 213–226 (1989). doi:10.1016/0306-4522(89)90120-6

[6] DeLong, M.R.: Primate models of movement disorders of basal ganglia origin. Trends Neurosci. 13(7), 281–285 (1990). doi:10.1016/0166-2236(90)90110-v

[7] Bergman, H., Wichmann, T., Karmon, B., DeLong, M.R.: The primate subthalamic nucleus. II. Neuronal activity in the MPTP model of parkinsonism. J. Neurophysiol. 72(2), 507–520 (1994). doi:10.1152/jn.1994.72.2.507

[8] Bergman, H., Feingold, A., Nini, A., Raz, A., Slovin, H., Abeles, M., Vaadia, E.: Physiological aspects of information processing in the basal ganglia of normal and parkinsonian primates. Trends Neurosci. 21(1), 32–38 (1998). doi:10.1016/s0166-2236(97)01151-x

[9] Marsden, C.D., Parkes, J.D.: “On-off” effects in patients with Parkinson’s disease on chronic levodopa therapy. Lancet 307(7954), 292–296 (1976). doi:10.1016/s0140-6736(76)91416-1

[10] Schrag, A., Quinn, N.: Dyskinesias and motor fluctuations in Parkinson’s disease: a community-based study. Brain 123(11), 2297–2305 (2000). doi:10.1093/brain/123.11.2297

[11] Shaw, K.M., Lees, A.J., Stern, G.M.: The impact of treatment with levodopa on Parkinson’s disease. QJM 49(3), 283–293 (1980). doi:10.1093/oxfordjournals.qjmed.a067623

[12] National Institute of Neurological Disorders and Stroke: Parkinson’s Disease. Available at https://www.ninds.nih.gov/health-information/disorders/parkinsons-disease. Accessed on 2025-11-28

[13] Schuurman, P.R., Bosch, D.A., Bossuyt, P.M.M., Bonsel, G.J., van Someren, E.J.W., de Bie, R.M.A., Merkus, M.P., Speelman, J.D.: A comparison of continuous thalamic stimulation and thalamotomy for suppression of severe tremor. N. Engl. J. Med. 342(7), 461–468 (2000). doi:10.1056/nejm200002173420703

[14] Wielepp, J.P., Burgunder, J.-M., Pohle, T., Ritter, E.P., Kinser, J.A., Krauss, J.K.: Deactivation of thalamocortical activity is responsible for suppression of parkinsonian tremor by thalamic stimulation: a 99mTc-ECD SPECT study. Clin. Neurol. Neurosurg. 103(4), 228–231 (2001). doi:10.1016/s0303-8467(01)00165-2

[15] Benabid, A.L., Pollak, P., Gervason, C., Hoffmann, D., Gao, D.M., Hommel, M., Perret, J.E., de Rougemont, J.: Long-term suppression of tremor by chronic stimulation of the ventral intermediate thalamic nucleus. Lancet 337(8738), 403–406 (1991). doi:10.1016/0140-6736(91)91175-t

[16] Alkemade, A., Forstmann, B.U.: Imaging of the human subthalamic nucleus. In: Swaab, D.F., Kreier, F., Lucassen, P.J., Salehi, A., Buijs, R.M. (eds.) The Human Hypothalamus: Middle and Posterior Region. Handbook of Clinical Neurology, vol. 180, pp. 403–416. Elsevier, Amsterdam, Netherlands (2021). Chap. 25. doi:10.1016/b978-0-12-820107-7.00025-2

[17] Limousin, P., Krack, P., Pollak, P., Benazzouz, A., Ardouin, C., Hoffmann, D., Benabid, A.-L.: Electrical stimulation of the subthalamic nucleus in advanced Parkinson’s disease. N. Engl. J. Med. 339(16), 1105–1111 (1998). doi:10.1056/nejm199810153391603

[18] Rodriguez-Oroz, M.C., Obeso, J.A., Lang, A.E., Houeto, J.-L., Pollak, P., Rehncrona, S., Kulisevsky, J., Albanese, A., Volkmann, J., Hariz, M.I., Quinn, N.P., Speelman, J.D., Guridi, J., Zamarbide, I., Gironell, A., Molet, J., Pascual-Sedano, B., Pidoux, B., Bonnet, A.M., Agid, Y., Xie, J., Benabid, A.-L., Lozano, A.M., Saint-Cyr, J., Romito, L., Contarino, M.F., Scerrati, M., Fraix, V., Van Blercom, N.: Bilateral deep brain stimulation in Parkinson’s disease: a multicentre study with 4 years follow-up. Brain 128(10), 2240–2249 (2005). doi:10.1093/brain/awh571

[19] Lanciego, J.L., Luquin, N., Obeso, J.A.: Functional neuroanatomy of the basal ganglia. Cold Spring Harb. Perspect. Med. 2, a009621 (2012). doi:10.1101/cshperspect.a009621

[20] Bejjani, B.-P., Gervais, D., Arnulf, I., Papadopoulos, S., Demeret, S., Bonnet, A.-M., Cornu, P., Damier, P., Agid, Y.: Axial parkinsonian symptoms can be improved: the role of levodopa and bilateral subthalamic stimulation. J. Neurol. Neurosurg. Psychiatry 68(5), 595–600 (2000). doi:10.1136/jnnp.68.5.595

[21] Dick, J.P.R.: Multicentre European study of thalamic stimulation in essential tremor. J. Neurol. Neurosurg. Psychiatry 74(10), 1362–1363 (2003). doi:10.1136/jnnp.74.10.1362

[22] Deuschl, G., Schade-Brittinger, C., Krack, P., Volkmann, J., Schäfer, H., Bötzel, K., Daniels, C., Deutschländer, A., Dillmann, U., Eisner, W., Gruber, D., Hamel, W., Herzog, J., Hilker, R., Klebe, S., Kloß, M., Koy, J., Krause, M., Kupsch, A., Lorenz, D., Lorenzl, S., Mehdorn, H.M., Moringlane, J.R., Oertel, W., Pinsker, M.O., Reichmann, H., Reuß, A., Schneider, G.-H., Schnitzler, A., Steude, U., Sturm, V., Timmermann, L., Tronnier, V., Trottenberg, T., Wojtecki, L., Wolf, E., Poewe, W., Voges, J.: A randomized trial of deep-brain stimulation for Parkinson’s disease. N. Engl. J. Med. 355(9), 896–908 (2006). doi:10.1056/nejmoa060281

[23] Chandra, V., Hilliard, J.D., Foote, K.D.: Deep brain stimulation for the treatment of tremor. J. Neurol. Sci. 435, 120190 (2022). doi:10.1016/j.jns.2022.120190

[24] Temperli, P., Ghika, J., Villemure, J.-G., Burkhard, P.R., Bogousslavsky, J., Vingerhoets, F.J.G.: How do parkinsonian signs return after discontinuation of subthalamic DBS? Neurology 60(1), 78–81 (2003). doi:10.1212/wnl.60.1.78

[25] Kromer, J.A., Khaledi-Nasab, A., Tass, P.A.: Impact of number of stimulation sites on long-lasting desynchronization effects of coordinated reset stimulation. Chaos 30(8), 083134 (2020). doi:10.1063/5.0015196

[26] Voges, J., Volkmann, J., Allert, N., Lehrke, R., Koulousakis, A., Freund, H.-J., Sturm, V.: Bilateral high-frequency stimulation in the subthalamic nucleus for the treatment of Parkinson disease: correlation of therapeutic effect with anatomical electrode position. J. Neurosurg. 96(2), 269–279 (2002). doi:10.3171/jns.2002.96.2.0269

[27] Tass, P.A.: A model of desynchronizing deep brain stimulation with a demand-controlled coordinated reset of neural subpopulations. Biol. Cybern. 89(2), 81–88 (2003). doi:10.1007/s00422-003-0425-7

[28] Bin-Mahfoodh, M., Hamani, C., Sime, E., Lozano, A.M.: Longevity of batteries in internal pulse generators used for deep brain stimulation. Stereotact. Funct. Neurosurg. 80(1–4), 56–60 (2004). doi:10.1159/000075161

[29] Tass, P.A.: Desynchronization by means of a coordinated reset of neural sub-populations: a novel technique for demand-controlled deep brain stimulation. Prog. Theor. Phys. Suppl. 150, 281–296 (2003). doi:10.1143/ptps.150.281

[30] Adamchic, I., Hauptmann, C., Barnikol, U.B., Pawelczyk, N., Popovych, O., Barnikol, T.T., Silchenko, A., Volkmann, J., Deuschl, G., Meissner, W.G., Maarouf, M., Sturm, V., Freund, H.-J., Tass, P.A.: Coordinated reset neuromodulation for Parkinson’s disease: proof-of-concept study. Mov. Disord. 29(13), 1679–1684 (2014). doi:10.1002/mds.25923

[31] Tass, P.A.: Effective desynchronization by means of double-pulse phase resetting. Europhys. Lett. 53(1), 15–21 (2001). doi:10.1209/epl/i2001-00117-6

[32] Tass, P.A.: Phase Resetting in Medicine and Biology: Stochastic Modelling and Data Analysis. Springer, Berlin, Germany (1999)

[33] Tass, P.A.: Desynchronizing double-pulse phase resetting and application to deep brain stimulation. Biol. Cybern. 85(5), 343–354 (2001). doi:10.1007/s004220100268

[34] Tass, P.A.: Desynchronization of brain rhythms with soft phase-resetting techniques. Biol. Cybern. 87(2), 102–115 (2002). doi:10.1007/s00422-002-0322-5

[35] Wang, X., Yu, Y., Wang, Q.: Modulating propagation of parkinsonian beta oscillations with transcranial direct current stimulation: a computational study. Chaos Solitons Fractals 189(1), 115635 (2024). doi:10.1016/j.chaos.2024.115635

[36] Yu, Y., Wang, H., Liu, X., Wang, Q.: Closed-loop transcranial electrical stimulation for inhibiting epileptic activity propagation: a whole-brain model study. Nonlinear Dyn. 112(23), 21369–21387 (2024). doi:10.1007/s11071-024-10132-w

[37] Shen, K.-Z., Zhu, Z.-T., Munhall, A., Johnson, S.W.: Synaptic plasticity in rat subthalamic nucleus induced by high-frequency stimulation. Synapse 50(4), 314–319 (2003). doi:10.1002/syn.10274

[38] Caporale, N., Dan, Y.: Spike timing–dependent plasticity: a Hebbian learning rule. Annu. Rev. Neurosci. 31, 25–46 (2008). doi:10.1146/annurev.neuro.31.060407.125639

[39] Ebert, M., Hauptmann, C., Tass, P.A.: Coordinated reset stimulation in a largescale model of the STN-GPe circuit. Front. Comput. Neurosci. 8, 154 (2014). doi:10.3389/fncom.2014.00154

[40] Khaledi-Nasab, A., Kromer, J.A., Tass, P.A.: Long-lasting desynchronization of plastic neuronal networks by double-random coordinated reset stimulation. Front. Netw. Physiol. 2, 864859 (2022). doi:10.3389/fnetp.2022.864859

[41] Tass, P.A., Majtanik, M.: Long-term anti-kindling effects of desynchronizing brain stimulation: a theoretical study. Biol. Cybern. 94(1), 58–66 (2006). doi:10.1007/s00422-005-0028-6

[42] Fan, D., Wang, Q.: Improving desynchronization of parkinsonian neuronal network via triplet-structure coordinated reset stimulation. J. Theor. Biol. 370, 157–170 (2015). doi:10.1016/j.jtbi.2015.01.040

[43] Manos, T., Zeitler, M., Tass, P.A.: How stimulation frequency and intensity impact on the long-lasting effects of coordinated reset stimulation. PLoS Comput. Biol. 14(5), e1006113 (2018). doi:10.1371/journal.pcbi.1006113

[44] Izhikevich, E.M.: Which model to use for cortical spiking neurons? IEEE Trans. Neural Netw. 15(5), 1063–1070 (2004). doi:10.1109/tnn.2004.832719

[45] Rubin, J.E., Terman, D.: High frequency stimulation of the subthalamic nucleus eliminates pathological thalamic rhythmicity in a computational model. J. Comput. Neurosci. 16(3), 211–235 (2004). doi:10.1023/b:jcns.0000025686.47117.67

[46] Kromer, J.A., Tass, P.A.: Coordinated reset stimulation of plastic neural networks with spatially dependent synaptic connections. Front. Netw. Physiol. 4, 1351815 (2024). doi:10.3389/fnetp.2024.1351815

[47] Kubota, S., Rubin, J.E.: Numerical optimization of coordinated reset stimulation for desynchronizing neuronal network dynamics. J. Comput. Neurosci. 45(1), 2018 (45–58). doi:10.1007/s10827-018-0690-z

[48] Liu, C., Yao, Y., Wang, J., Li, H., Wu, H., Loparo, K.A., Fietkiewicz, C.: Oscillation suppression effects of intermittent noisy deep brain stimulation induced by coordinated reset pattern based on a computational model. Biomed. Signal Process. Control 73, 103466 (2022). doi:10.1016/j.bspc.2021.103466

[49] Hindmarsh, J.L., Rose, R.M.: A model of neuronal bursting using three coupled first order differential equations. Proc. R. Soc. Lond. B Biol. Sci. 221(1222), 87–102 (1984). doi:10.1098/rspb.1984.0024

[50] Malik, S.A., Mir, A.H.: Synchronization of Hindmarsh Rose neurons. Neural Netw. 123, 372–380 (2020). doi:10.1016/j.neunet.2019.11.024

[51] Jalili, M.: Phase synchronizing in Hindmarsh–Rose neural networks with delayed chemical coupling. Neurocomputing 74(10), 1551–1556 (2011). doi:10.1016/j.neucom.2010.12.031

[52] Jalili, M.: Collective behavior of interacting locally synchronized oscillations in neuronal networks. Commun. Nonlinear Sci. Numer. Simul. 17(10), 3922–3933 (2012). doi:10.1016/j.cnsns.2012.02.005

[53] Chakravartula, S., Indic, P., Sundaram, B., Killingback, T.: Emergence of local synchronization in neuronal networks with adaptive couplings. PLoS One 12(6), e0178975 (2017). doi:10.1371/journal.pone.0178975

[54] Baran, A.Y., Korkmaz, N., Öztürk, I., Kılıç, R.: On addressing the similarities between STDP concept and synaptic/memristive coupled neurons by realizing of the memristive synapse based HR neurons. Eng. Sci. Technol. Int. J. 32, 101062 (2022). doi:10.1016/j.jestch.2021.09.008

[55] Lysyansky, B., Popovych, O.V., Tass, P.A.: Multi-frequency activation of neuronal networks by coordinated reset stimulation. Interface Focus 1(1), 75–85 (2011). doi:10.1098/rsfs.2010.0010

[56] Atwood, H.L., Wojtowicz, J.M.: Silent synapses in neural plasticity: current evidence. Learn. Mem. 6(6), 542–571 (1999). doi:10.1101/lm.6.6.542

[57] Bevan, M.D., Wilson, C.J.: Mechanisms underlying spontaneous oscillation and rhythmic firing in rat subthalamic neurons. J. Neurosci. 19(17), 7617–7628 (1999). doi:10.1523/jneurosci.19-17-07617.1999

[58] Hammond, C., Bergman, H., Brown, P.: Pathological synchronization in Parkinson’s disease: networks, models and treatments. Trends Neurosci. 30(7), 357–364 (2007). doi:10.1016/j.tins.2007.05.004

[59] Chen, D., Li, J., Zeng, W., He, J.: Topology identification and dynamical pattern recognition for Hindmarsh–Rose neuron model via deterministic learning. Cogn. Neurodyn. 17(1), 203–220 (2023). doi:10.1007/s11571-022-09812-3

[60] Heeger, D.: Poisson model of spike generation. Technical report, Stanford University (2000)

[61] Lévesque, J.-C., Parent, A.: GABAergic interneurons in human subthalamic nucleus. Mov. Disord. 20(5), 574–584 (2005). doi:10.1002/mds.20374

[62] Feher, J.: Quantitative Human Physiology: An Introduction, 2nd edn. Elsevier, London, United Kingdom (2017)

[63] Emmi, A., Antonini, A., Macchi, V., Porzionato, A., De Caro, R.: Anatomy and connectivity of the subthalamic nucleus in humans and non-human primates. Front. Neuroanat. 14, 13 (2020). doi:10.3389/fnana.2020.00013

[64] Somers, D., Kopell, N.: Rapid synchronization through fast threshold modulation. Biol. Cybern. 68(5), 393–407 (1993). doi:10.1007/bf00198772

[65] Remi, T., Subha, P.A.: Chemically coupled Hindmarsh–Rose neurons with cross interactions between membrane potential and magnetic flux. J. Phys. A Math. Theor. 56(34), 345701 (2023). doi:10.1088/1751-8121/ace56f

[66] Kita, H., Chang, H.T., Kitai, S.T.: The morphology of intracellularly labeled rat subthalamic neurons: a light microscopic analysis. J. Comp. Neurol. 215(3), 245–257 (1983). doi:10.1002/cne.902150302

[67] Gillies, A.J., Willshaw, D.J.: A massively connected subthalamic nucleus leads to the generation of widespread pulses. Proc. R. Soc. Lond. B Biol. Sci. 265(1410), 2101–2109 (1998). doi:10.1098/rspb.1998.0546

[68] Goldberg, D.E., Deb, K.: A comparative analysis of selection schemes used in genetic algorithms. Found. Genet. Algorithms 1, 69–93 (1991). doi:10.1016/b978-0-08-050684-5.50008-2

[69] Bevan, M.D., Magill, P.J., Terman, D., Bolam, J.P., Wilson, C.J.: Move to the 27 rhythm: oscillations in the subthalamic nucleus–external globus pallidus network. Trends Neurosci. 25(10), 525–531 (2002). doi:10.1016/s0166-2236(02)02235-x

[70] Park, C., Worth, R.M., Rubchinsky, L.L.: Fine temporal structure of beta oscillations synchronization in subthalamic nucleus in Parkinson’s disease. J. Neurophysiol. 103(5), 2707–2716 (2010). doi:10.1152/jn.00724.2009

[71] Rosenblum, M., Pikovsky, A., Kurths, J., Schäfer, C., Tass, P.A.: Phase synchronization: from theory to data analysis. In: Moss, F., Gielen, S. (eds.) Neuro-Informatics and Neural Modelling. Handbook of Biological Physics, vol. 4, pp. 279–321. Elsevier, Amsterdam, Netherlands (2001). Chap. 9. doi:10.1016/s1383-8121(01)80012-9

[72] Zeitler, M., Tass, P.A.: Computationally developed sham stimulation protocol for multichannel desynchronizing stimulation. Front. Physiol. 9, 512 (2018). doi:10.3389/fphys.2018.00512

[73] Lysyansky, B., Popovych, O.V., Tass, P.A.: Desynchronizing anti-resonance effect of m : n on–off coordinated reset stimulation. J. Neural Eng. 8(3), 036019 (2011). doi:10.1088/1741-2560/8/3/036019

[74] Zeitler, M., Tass, P.A.: Augmented brain function by coordinated reset stimulation with slowly varying sequences. Front. Syst. Neurosci. 9, 49 (2015). doi:10.3389/fnsys.2015.00049

[75] Perlmutter, J.S., Mink, J.W.: Deep brain stimulation. Annu. Rev. Neurosci. 29, 229–257 (2006). doi:10.1146/annurev.neuro.29.051605.112824

[76] Harnack, D., Winter, C., Meissner, W., Reum, T., Kupsch, A., Morgenstern, R.: The effects of electrode material, charge density and stimulation duration on the safety of high-frequency stimulation of the subthalamic nucleus in rats. J. Neurosci. Methods 138(1–2), 207–216 (2004). doi:10.1016/j.jneumeth.2004.04.019

[77] Sadeh, S., Clopath, C.: Excitatory-inhibitory balance modulates the formation and dynamics of neuronal assemblies in cortical networks. Sci. Adv. 7(45), eabg8411 (2021). doi:10.1126/sciadv.abg8411

[78] Liu, R., Zhang, Y., Wang, J., Zheng, M., Xu, K.: Formation and maintenance of neuronal collective dynamics through local perturbation and intrinsic node dynamics. Chaos Solitons Fractals 199(1), 116648 (2025). doi:10.1016/j.chaos.2025.116648

[79] Rappel, P., Grosberg, S., Arkadir, D., Linetsky, E., Abu Snineh, M., Bick, A.S., Tamir, I., Valsky, D., Marmor, O., Abo Foul, Y., Peled, O., Gilad, M., Daudi, C., Ben-Naim, S., Bergman, H., Israel, Z., Eitan, R.: Theta-alpha oscillations characterize emotional subregion in the human ventral subthalamic nucleus. Mov. Disord. 35(2), 337–343 (2020). doi:10.1002/mds.27910

[80] Yin, Z., Zhu, G., Zhao, B., Bai, Y., Jiang, Y., Neumann, W.-J., Kühn, A.A., Zhang, J.: Local field potentials in Parkinson’s disease: a frequency-based review. Neurobiol. Dis. 155, 105372 (2021). doi:10.1016/j.nbd.2021.105372

[81] Bi, G.-Q., Poo, M.-M.: Synaptic modifications in cultured hippocampal neurons: dependence on spike timing, synaptic strength, and postsynaptic cell type. J. Neurosci. 18(24), 10464–10472 (1998). doi:10.1523/jneurosci.18-24-10464.1998

[82] Asl, M.M., Vahabie, A.-H., Valizadeh, A., Tass, P.A.: Spike-timing-dependent plasticity mediated by dopamine and its role in Parkinson’s disease pathophysiology. Front. Netw. Physiol. 2, 817524 (2022). doi:10.3389/fnetp.2022.817524

[83] Chauhan, K., Neiman, A.B., Tass, P.A.: Synaptic reorganization of synchronized neuronal networks with synaptic weight and structural plasticity. PLoS Comput. Biol. 20(7), e1012261 (2024). doi:10.1371/journal.pcbi.1012261

[84] Reck, C., Florin, E., Wojtecki, L., Krause, H., Groiss, S., Voges, J., Maarouf, M., Sturm, V., Schnitzler, A., Timmermann, L.: Characterisation of tremor-associated local field potentials in the subthalamic nucleus in Parkinson’s disease.Eur. J. Neurosci. 29(3), 599–612 (2009). doi:10.1111/j.1460-9568.2008.06597.x

[85] Beudel, M., Little, S., Pogosyan, A., Ashkan, K., Foltynie, T., Limousin, P., Zrinzo, L., Hariz, M., Bogdanovic, M., Cheeran, B., Green, A.L., Aziz, T., Thevathasan, W., Brown, P.: Tremor reduction by deep brain stimulation is associated with gamma power suppression in Parkinson’s disease. Neuromodulation 18(5), 349–354 (2015). doi:10.1111/ner.12297

[86] Liu, C., Wang, J., Li, H., Fietkiewicz, C., Loparo, K.A.: Modeling and analysis of beta oscillations in the basal ganglia. IEEE Trans. Neural Netw. Learn. Syst. 29(5), 1864–1875 (2018). doi:10.1109/tnnls.2017.2688426

