## Supplementary Material for "Hindmarsh–Rose neuronal network with spike-timing-dependent plasticity demonstrates coordinated reset neuromodulation for Parkinson’s disease"

Shahin Sharafi, Jesse I. Gilmer, Mazen Al Borno, Thomas K. Uchida

### A Examples of Noise Used in Hindmarsh–Rose Model

As discussed in Sect. 2.1, two noise models were explored in this work. The first is simple white noise, where a random value is selected at each time step from a uniform distribution over the range $[-0.5, 0.5]$. This noise model assumes that the input received from all unmodelled neurons, once these inputs have been scaled by their respective synaptic weights and summed, resembles white noise. The second noise model assumes the unmodelled neurons provide an input that resembles an independent Poisson process, with firing rate 50 Hz, noise intensity 3.0, and synaptic timescale 2 ms. Examples of these noise inputs $i^{\text{noise}}(t)$ are illustrated in Supp. Figs. 1 and 2.

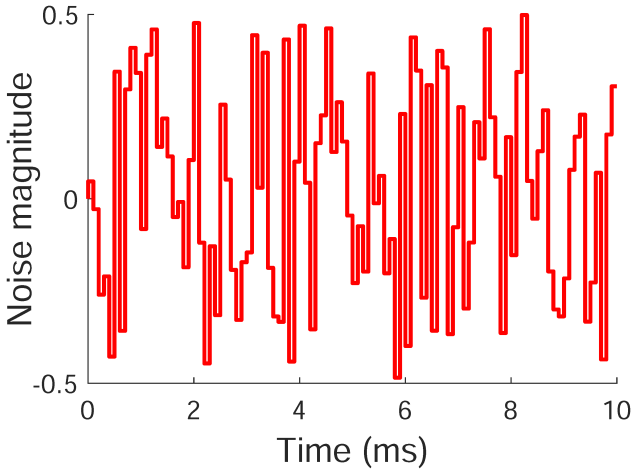


**Supplementary Figure 1:** Example of uniformly distributed white noise.

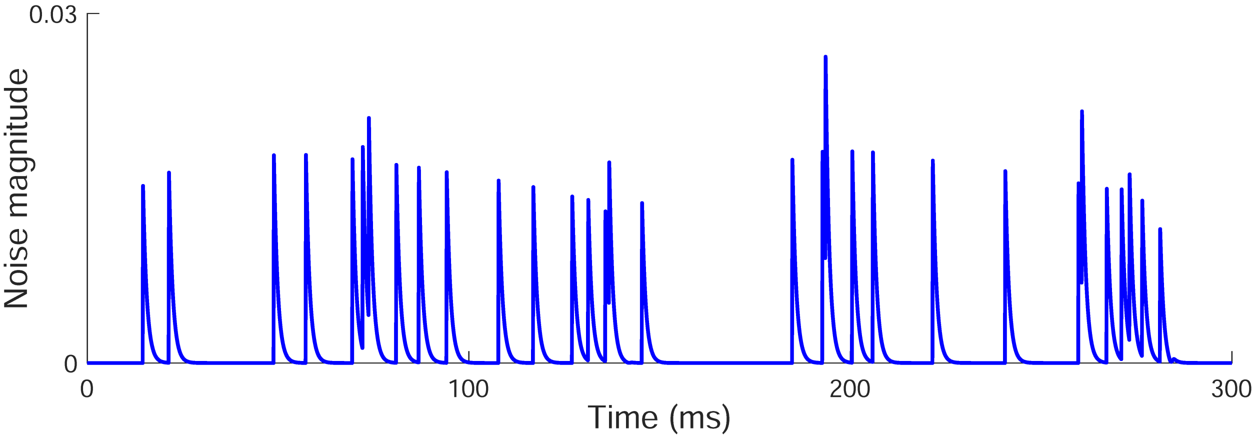


**Supplementary Figure 2:** Example of Poisson noise.
